# The AP-1 factors *FOSL1* and *FOSL2* co-regulate human Th17 responses

**DOI:** 10.1101/2021.04.26.441472

**Authors:** Ankitha Shetty, Subhash Kumar Tripathi, Sini Junttila, Tanja Buchacher, Rahul Biradar, Santosh D. Bhosale, Tapio Envall, Asta Laiho, Robert Moulder, Omid Rasool, Sanjeev Galande, Laura L. Elo, Riitta Lahesmaa

**Affiliations:** Turku Bioscience Centre, University of Turku and Åbo Akademi University; Turku 20520, Finland; InFLAMES Research Flagship Center, University of Turku; Centre of Excellence in Epigenetics, Department of Biology, Indian Institute of Science Education and Research (IISER); Pune 411008, India; The Leibniz Institute for Natural Product Research and Infection Biology – Hans Knöll Institute; Beutenbergstraße 13, 07745 Jena, Germany; Department of Biochemistry and Molecular Biology, Protein Research Group, University of Southern Denmark; Campusvej 55, Odense M, DK-5230, Denmark

## Abstract

Th17 cells protect mucosal barriers, but their aberrant activity can cause autoimmunity. Molecular networks dictating human Th17 function are largely unexplored, and this hinders disease-studies. Here, we investigated the roles of the AP-1 factors, *FOSL1* and *FOSL2,* in inducing human Th17 responses. Transient knockdown and over-expression strategies found the two proteins to inhibit Th17-cell identity, while revealing a distinct cooperativity between their functions. Strikingly, *FOSL1* plays different roles in human and mouse and FOSL-mediated Th17 regulation is opposed by the AP-1 factor, BATF. Genome-wide occupancy analysis demonstrated the co-localization of FOSL1, FOSL2 and BATF in the vicinity of key Th17 genes. The functional interplay among these factors is possibly governed by sharing interactions with a common set of lineage-associated proteins. We further discovered that the genomic binding sites of these factors harbour a large number of disease-linked SNPs, many of which alter the ability of a given factor to bind DNA. Our findings thus provide crucial insights into the transcriptional regulation of human Th17 function and associated pathologies.

**ONE SENTENCE SUMMARY:** FOSL1- and FOSL2-mediated transcription during early human Th17 differentiation

## INTRODUCTION

Th17 cells are crucial for mucosal defence against extracellular bacteria and fungi. Their uncontrolled response, however, can lead to autoimmune conditions such as rheumatoid arthritis (RA), multiple sclerosis (MS) and inflammatory bowel disease (IBD). Characterization of the molecular circuits that dictate Th17 cell-function, is thus critical for the therapeutic development of immune-mediated disorders. Primarily, Th17 differentiation is initiated when naive CD4^+^ T cells are exposed to IL-6 and TGF-β (with or without IL-1β or IL-23). The early stages of differentiation involve signaling cascades that endorse lineage-defining gene-expression programs and restrict the diversification to other T-helper cell fates. These events are dictated by a well-coordinated network of transcription factors (TFs), many of which have been functionally characterized in both human and mouse. Key studies using gene-knockout mouse models have demonstrated how pioneer factors, such as STAT3, BATF and IRF4, nucleate key Th17-defining proteins (RORγT, RORα) over cytokine gene loci (*IL-17A, IL17F* and *IL22)* (*1*).

The AP-1 complex, which is comprised of JUN, FOS and ATF family members, is known to control Th17-regulatory circuits in mouse (*1–6*). Included within the FOS family are *FOSL1* and *FOSL2* (also known as *FRA-1* or *FRA-2*), two paralogous TFs that have limited sequence similarity (< 40%) and can perform different functions. Their importance in the regulation of embryonic development, cancer progression, ECM synthesis and immune cell responses is well-established (*7–13*). In addition, they are known to exert opposite effects on murine Th17-development. While *FOSL1* positively regulates the process, *FOSL2* acts as a repressor of the lineage (*1, 3, 4*). Though these molecular players have been prodigiously studied in mouse, their roles have not been confirmed in human. Heterogeneity between human and mouse with regard to cytokine-responses, T cell-activation, γ/δ T-cell function and interferon signalling, are quite well-known (*14*). Significant differences have also been reported for the proteomic profiles of early-differentiating Th17 cells in the two species (*15*). Moreover, genes such as *AHR* (*16, 17*), *PRDM1* (*18, 19*) and *SATB1* (*15*), have been found to exhibit divergent functions in human and mouse Th17-regulation. These discrepancies should be especially borne in mind while extrapolating information from murine studies for therapeutic interventions. They further underscore the need for validating murine gene-functions using human cells.

FOS and ATF proteins lack a transactivation domain, and thus need to heterodimerize with JUN and other factors to execute their gene-regulatory roles (*8, 20, 21*). Since the resulting transcriptional activity is dictated by both of the proteins forming the dimer, dissecting the individual function of the monomers has proved to be challenging. Findings across cell types have identified both co-operative and antagonistic relationships among AP-1 proteins, in a context-specific manner (*8, 21*). Thus, investigating the AP-1 complex requires more comprehensive approaches where the molecular interplay between its members could be addressed. Such interconnectivity is previously reported for FOSL2 and BATF in murine Th17 cells, where the two factors share occupancy over lineage-specific gene loci, yet regulate Th17-fate in an opposite fashion (*1, 5*). While their interplaying roles are largely unexplored in human T cells, the functional relationship between FOSL1 and BATF stands undetermined in either of the species.

In the present study, we investigated the individual and connected roles of FOSL1, FOSL2 and BATF in regulating human Th17 cell-identity, while highlighting species-specific differences. By combining global gene-expression analysis and genome-wide occupancy studies, we dissected the genes that are directly-regulated by these TFs. Our results demonstrate an evident cooperation between FOSL1 and FOSL2 functions, while verifying their antagonistic relationship with BATF in human Th17 cells. Further analysis revealed that the genomic regions bound by these AP-1 proteins harbour hundreds of disease-linked single nucleotide polymorphisms (SNPs), many of which altered the ability of these proteins to bind DNA. Disrupting the binding-affinities of these TFs to their target gene-regulatory sites could subsequently alter their roles in instrumenting Th17 responses, and thereby contribute to disease development. Together, findings from this study provide a better understanding of the transcriptional regulation of human Th17 lineage, which could help in designing new therapeutic strategies for associated diseases.

## RESULTS

### FOSL1 and FOSL2 negatively influence early human Th17 responses

The Th17-specific expression profile of FOS-like proteins was examined using RNA-seq data from our previously published study (*22*). FOSL1 and FOSL2 transcript levels were plotted for umbilical cord blood-derived naive CD4^+^ T cells, which were cultured under activation (Th0) or Th17-polarizing conditions (TGF-β, IL-6 and IL-1β), for different time points (Fig. 1A). When compared to activated T cells, the Th17-lineage showed a significant increase in levels of both factors. These changes were subsequently validated at the protein level by immunoblot analysis (Fig. 1B; Fig S1A). While both proteins were the most differentially upregulated at 24h, FOSL2 depicted a more-striking trend at all of the evaluated time points.

**Fig. 1.**
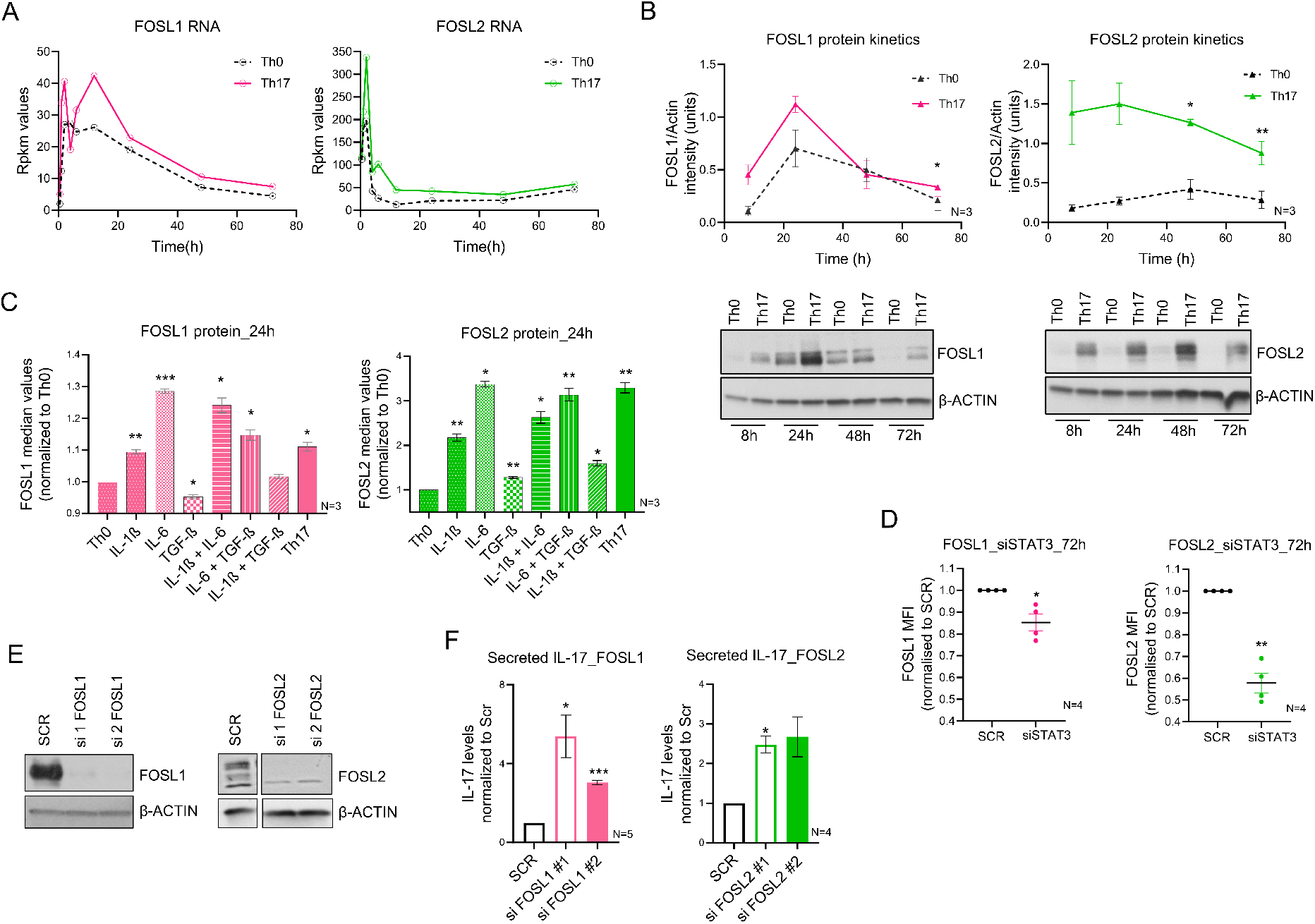
Expression and function of FOS-like proteins during human Th17 differentiation. (**A**) Rpkm values are plotted for FOSL1 [left] and FOSL2 [right] RNA at different time points of activation (Th0) or Th17-polarization, using published RNA-seq data (*Tuomela et al., 2016 Oncotarget*). (**B**) Immunoblot images (lower panel) show FOSL1 [left] and FOSL2 [right] protein levels in Th0 and Th17-polarizing cells, over a time-course. Actin was used as loading control. Blots from three biological replicates were quantified using ImageJ and the corresponding FOSL intensity values (normalized to actin) are plotted as a line graph in the above panel. (**C**) Flow cytometry analysis of FOSL1 [left] and FOSL2 [right] expression in naive CD4+ T cells cultured for 24h, under conditions of activation (Th0), Th17-polarization, or activation in presence of the Th17-cytokines (used either alone or in combination). Bar plot shows median fluorescence intensity (MFI) values normalized to Th0, for three biological replicates. Statistical significance was calculated by comparing each condition to Th0. (**D**) Flow cytometry analysis of FOSL1 [left] and FOSL2 [right] protein levels in non-targeting (SCR) versus STAT3 KD Th17 cells, at 72h of polarization. Graph shows MFI values normalized to SCR for four biological replicates. (**E**) Naive CD4^+^ T cells were silenced for FOSL1 [left] or FOSL2 [right] using two different siRNAs each, and further polarized to Th17-fate for 24h. Knockdown was analyzed using immunoblotting. Representative blots for three biological replicates are shown. (**F**) ELISA was used to estimate IL-17A secretion in supernatants of FOSL1 [left] and FOSL2-silenced [right] Th17 cells, at 72h of polarization. Values were first normalized to live cell count, followed by normalization with SCR. Data represents four or five biological replicates, as indicated. Graphs in the above panels show mean ± standard error of the mean (SEM). Statistical significance is calculated using two-tailed Student’s t test (*p < 0.05; **p < 0.01, ***p < 0.001).

TCR signaling is known to upregulate AP-1 activity (*8, 23*). We thus investigated which of the Th17-polarizing cytokines increase FOSL expression above TCR-induced levels. To achieve this, naive CD4^+^ T cells were activated in the presence of different Th17 cytokines, used either individually or in combination. FOSL1 and FOSL2 protein levels were then analysed at 24h of polarization, using flow cytometry (Fig. 1C). Our results found IL-1β and IL-6 to significantly enhance the expression of both proteins (relative to Th0); however, IL-6 showed a stronger effect. In addition, TGF-β suppressed FOSL1 expression but promoted FOSL2 levels, which complies with previous findings (*24, 25*).

The IL-6/STAT3 signaling axis is known to drive the expression of FOS-like proteins (*3, 15, 26–28*). Given the importance of STAT3 in establishing Th17-cell identity (*26, 29*), we sought to determine if the IL-6-induced increase in FOSL expression required STAT3 function in Th17 cells. To address this, FOSL1 and FOSL2 levels were examined in STAT3-depleted Th17-polarized cells, by immunoblotting (Fig. 1D). Although the loss of STAT3 reduced the expression of both factors, the effect was more pronounced on FOSL2. Notably, in an earlier study (*26*), we have found STAT3 to occupy the promoter region of *FOSL2* but not *FOSL1,* which might explain the more robust effect on the former (Fig. S1B).

The early and sustained expression of FOS-like proteins suggests their potential involvement in steering Th17-differentiation. To determine their precise roles, we silenced each of these proteins individually with RNAi and probed for an effect on IL-17 cytokine, which is a key marker of the Th17 lineage. To ensure reproducibility with minimal off-target phenotypes, FOSL1 and FOSL2 were each targeted using two different siRNAs. Naive CD4^+^ T cells were nucleofected and cultured according to the workflow in Fig. S1C. The siRNA-efficacy was confirmed with western blotting (Fig. 1E; Fig. S1D). Interestingly, depletion of FOSL1 or FOSL2 significantly increased IL-17 secretion at 72h of polarization, which highlights the negative influence of these factors on human Th17 cell-function (Fig. 1F). This also proves that although FOSL2 shows similar functions, FOSL1 exhibits divergent roles in human and mouse Th17 differentiation (*1, 3*).

### FOSL1 and FOSL2 synergistically repress IL-17 expression

FOS and JUN proteins can cooperatively regulate gene-expression (*30, 31*). Whether such synergy exists between FOSL1 and FOSL2 functions, is yet to be explored. Since both of them regulated IL-17 secretion in a similar fashion, we examined if their simultaneous perturbation causes enhanced changes. To achieve this, siRNA knockdown (KD) as well as RNA-based over-expression (OE) strategies were used. For simultaneous silencing (double KD or DKD), naive CD4^+^ T cells were nucleofected with a combination of FOSL1- and FOSL2-targeting siRNAs, and immunoblotting was performed to confirm the parallel reduction in their protein levels (Fig. 2A; Fig. S2A). Cells individually silenced for either of these factors served as single KD controls. In agreement with earlier findings (*32, 33*), silencing FOSL1 did not alter FOSL2 expression, and vice-versa (Fig. 2A). To further assess the effect of the KD on the expression of IL-17 cytokine, qPCR and ELISA analyses were performed at 72h of polarization. Notably, FOSL1 and FOSL2 co-depletion additively augmented IL-17 levels, relative to the single KD controls (Fig. 2C and E).

**Fig. 2.**
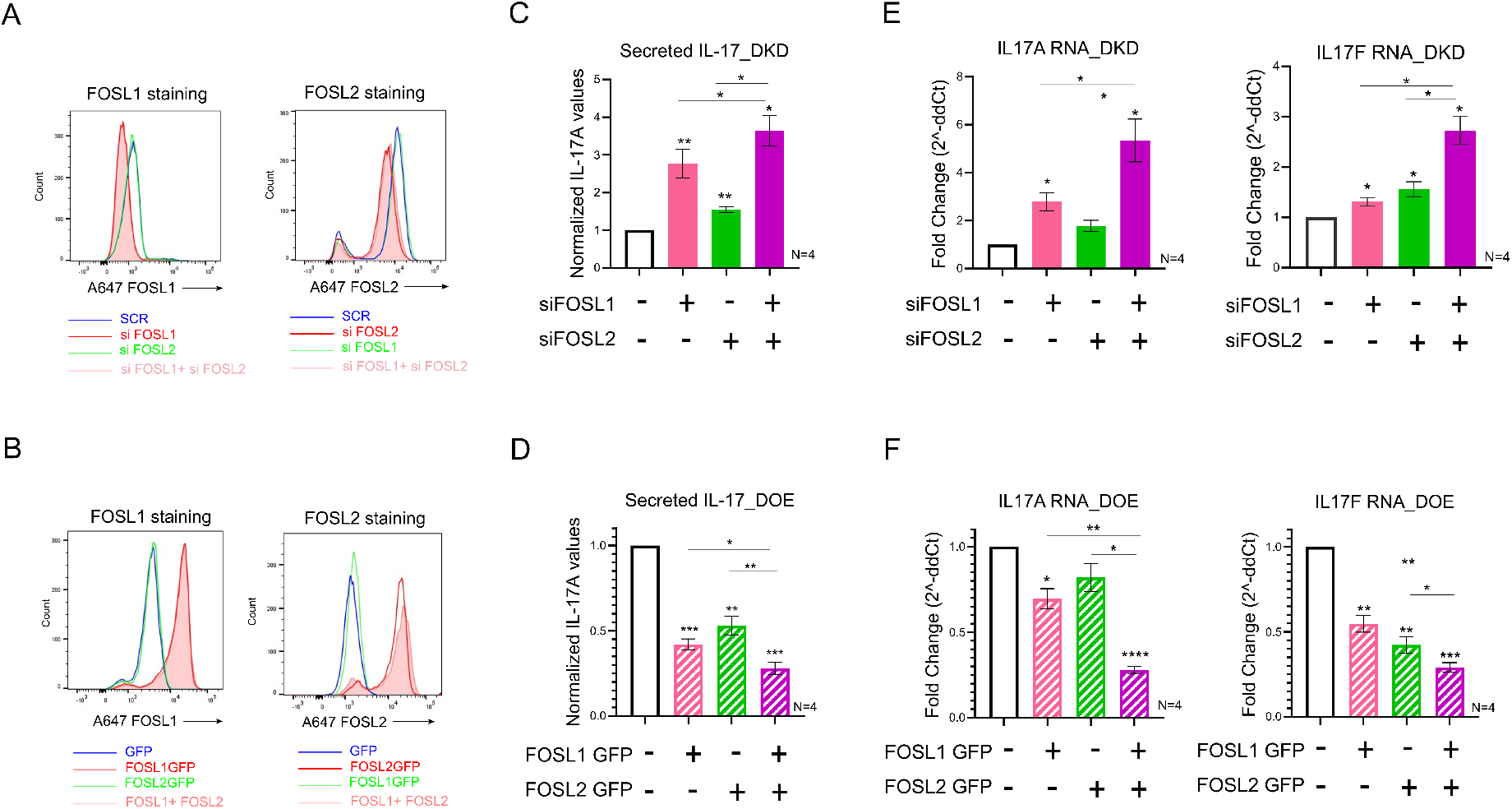
Cooperative regulation of IL-17 expression by FOSL1 and FOSL2. (**A**) Naive CD4^+^ T cells were silenced for FOSL1, FOSL2 or both factors in parallel (double KD; DKD), and cultured under Th17-polarizing conditions for 24h. Total FOSL1 [left] or FOSL2 [right] protein was stained (Alexa-647) and analysed by flow cytometry. Non-targeting siRNA (SCR) was used as nucleofection control. Representative histograms for four biological replicates are shown. (**B**) Naive CD4^+^ T cells were treated with *in-vitro* transcribed GFP-FOSL1 RNA, GFP-FOSL2 RNA or both (double OE; DOE). After resting the cells for 18–20h, total FOSL1 [left] or FOSL2 [right] protein was stained (Alexa-647) and analysed by flow cytometry. GFP RNA were used as nucleofection control. Representative histograms for four biological replicates are shown. (**C** and **D**) Bar plot shows ELISA results for secreted IL-17A levels in supernatants of FOSL KD/DKD (panel C) or FOSL OE/DOE Th17 cells (panel D), at 72h of polarization. Values were first normalized to live cell count, and then to the respective control condition (SCR or GFP). Data represent four biological replicates. (**E** and **F**) qRT-PCR analysis for measurement of IL-17A [left] and IL-17F [right] RNA levels in FOSL KD/DKD (panel E) or FOSL OE/DOE Th17 cells (panel F), at 72h of polarization. Fold-change normalized to the respective controls (SCR or empty GFP) was plotted for four biological replicates. For panels C-F, plots show mean ± SEM. Statistical significance is calculated using two-tailed Student’s t test (*p < 0.05, **p < 0.01, ***p < 0.001, ****p<0.0001).

To authenticate these findings, we simultaneously over-expressed the two proteins, using *in-vitro* transcribed (IVT) RNA. Naive CD4^+^ T cells were nucleofected with a combination of FOSL1 and FOSL2 IVT RNAs (double OE or DOE), and flow cytometry analysis confirmed the lateral increase in their levels (Fig. 2B; Fig. S2B). Parallel overexpression of the two proteins caused an additive inhibition in IL-17 levels, as compared to the single OE controls (Fig. 2D and F). This strengthened our RNAi findings and confirmed the functional coordination between FOSL1 and FOSL2.

### Perturbing FOS-like proteins triggers important changes in Th17 gene-expression programs

To globally unravel the individual and combined gene targets of FOSL proteins, RNA-sequencing and differential expression (DE) analysis was performed for KD and DKD Th17 cells. For FOSL1, FOSL2 and DKD conditions, respectively, our analysis detected 466, 1,529 and 2,000 DE genes at 24h and 315, 150 and 1,500 DE genes at 72h of polarization (false discovery rate (FDR) ≤ 0.1). A similar analysis was performed for OE and DOE Th17 cells (at 72h) resulting in the identification of 30, 352 and 522 DE transcripts for FOSL1, FOSL2 and DOE conditions, respectively (FDR ≤ 0.1). It was thus evident that co-perturbing these factors altered a higher number of genes.

To further identify the cooperatively-regulated targets, the fold-changes for the affected genes were compared in KD versus DKD and OE versus DOE conditions. A significant number of targets showed more pronounced expression changes when both the factors were simultaneously perturbed (Fig. 3A and B). These included key Th17-marker genes such as *IL17A, IL17F, IL23R and CCR6,* all of which were negatively-regulated. Similarly, other genes that control Th17-cell function and autoimmunity, such as *FASLG (34, 35), IL7R (36), NT5E (37–40), STAT4 (41–44), CD70 (45, 46), PRDM1 (19), FGF2 (47), DUSP2 (48), IL12RB1(49), IL11 (50–52), IL24 (53–55) and IRF7 (56)),* were also found to be co-regulated. Nonetheless, a small number of lineage-associated factors (*IL21, USP18, GZMB, IL3, and others)* were altered by FOSL1 and FOSL2 in a non-synergistic fashion. This suggests that apart from their coordinated roles, these proteins also independently guide Th17-gene networks.

**Fig. 3.**
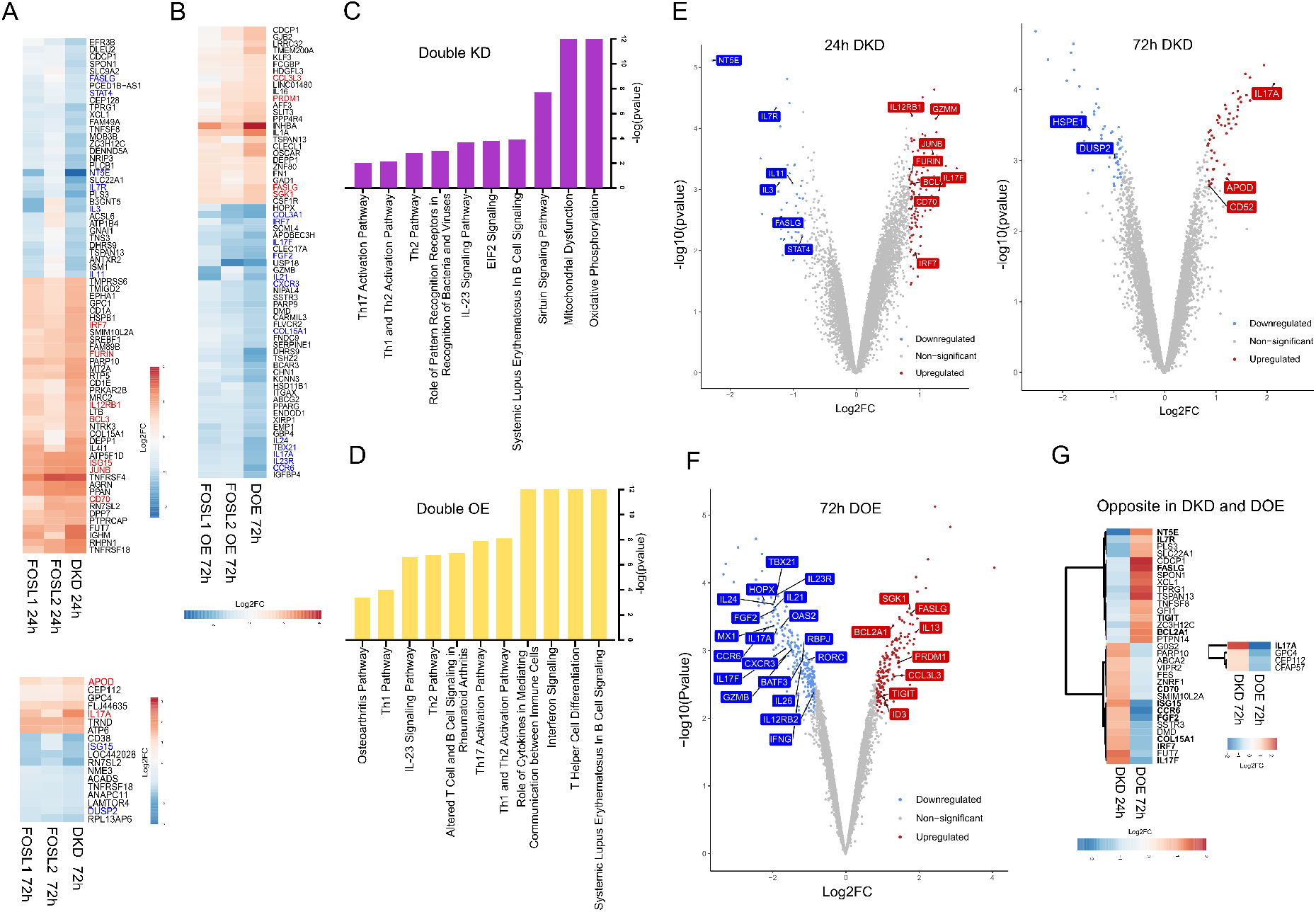
Transcriptional targets co-regulated by FOSL1 and FOSL2. (**A** and **B**) Heatmap in panel A shows the DE genes that are more profoundly altered in FOSL DKD Th17 cells relative to the single KD controls, at 24h (above) and 72h (below) of polarization. Panel B includes the DE genes that show enhanced changes in FOSL DOE Th17 cells as compared to the single OE controls at 72h of polarization. Genes with Th17-relevance are highlighted; upregulated genes are in red, and downregulated ones are in blue. Log2fold-change was calculated relative to the respective control conditions (i.e., SCR or GFP). (**C** and **D**) Ingenuity pathway analysis was used to identify signaling pathways that are altered upon FOSL DKD (panel C) or DOE (panel D). (**E** and **F)** Genome-wide expression analysis of FOSL DKD and FOSL DOE Th17 cells. Volcano plots in Panel E highlight the Th17-associated transcripts that are differentially expressed upon co-depletion of FOSL1 and FOSL2, at 24h [left] and 72h [right] of Th17 polarization. Panel F highlights the Th17-associated genes that are differentially expressed upon parallel over-expression of FOSL1 and FOSL2, at 72h of Th17 polarization. Targets with FDR ≤ 0.1 and |fold-change| ≥ 1.8 have been plotted. Upregulated genes are in red, and the downregulated ones are in blue. (**G**) Heatmap depicts the DE genes that show opposite expression changes in FOSL DKD versus DOE conditions, at the indicated time points of Th17 polarization. Th17-relevant genes have been highlighted.

Since co-perturbation affected a wider range of T-cell-related genes, we hereby focused on analysing the DKD and DOE datasets. In the case of DKD, our analysis revealed a poor overlap between the targets identified at 24h and 72h of polarization, which suggests time-distinguished functions (Fig. S3C). Ingenuity pathway analysis (IPA) further indicated that FOSL KD or OE perturbed genes involved mitochondrial dysfunction, oxidative phosphorylation, T-helper cell differentiation, IL-23 signaling, Interferon signaling, Th1/Th2/Th17 activation, and autoimmune-associated processes (RA, systemic lupus erythematosus (SLE)) (Fig. 3C and D).

The KD and OE data combined, interestingly revealed that FOSL proteins jointly repress several genes that characterize Th17 cell-identity (*RORC*, *FGF2 (47), IL21* (*57*)*, JUNB* (*2, 6*)*, CD70 (45, 46), IL12RB1* (*49*)*, CD52 (58, 59)* and *RBPJ (60))* and autoimmune phenotypes (*OAS2, MX1* and *ISG15* (*61*)) (Fig. 3E and F; Fig. S3A and B). At the same time, genes that inhibit Th17 cell-function and associated inflammation (*IL13* (*62*), *IL7R (36)*, *PRDM1 (19), DUSP2 (48), BCL2A1* (*63*), *ID3* (*64*)*, TIGIT (65)* and *NT5E (37–40)),* were found to be positively regulated by these factors (Fig. 3E and F; Fig. S3A and B). We also found known-targets of STAT3 (*HOPX, STAT4, IL23R, CCR6, IL24, GATA3, FNDC9, NR4A2, GZMB)* (*26*), which is a master-regulator of the Th17 lineage, to be influenced by FOSL in the opposite-direction. This ascertains the role of FOSL proteins in preventing the induction of Th17-fate.

To further identify the strongest synergistic targets of FOSL1 and FOSL2, the subset of genes that showed contrasting expression changes in DKD versus DOE conditions were selected (Fig. 3G). We found 37 such genes, many of which were functionally associated with the Th17 lineage. These included *IL17F, IL17A, CCR6 (66–68), FASLG (34, 35), FGF2 (47), IL7R (36), BCL2A1 (63), TIGIT (65), NT5E* (*37, 38, 40*)*, CD70 (45, 46)* and *IRF7 (56)*. Out of these, we validated the expression changes for CCR6 (*68*) which is a Th17-specific chemokine receptor, using flow cytometry analysis (Fig. S3D). To additionally authenticate our RNA-seq findings, the cooperative effects of FOSL were confirmed on other lineage-associated targets (*NT5E, STAT4, CD70, APOD, JUNB*), using either immunoblotting or flow cytometry (Fig. S3E; Fig. S4A-D). NT5E or CD73 is a 5′ectonucleotidase, which is known to resolve uncontrolled inflammation (*38, 69*). A positive correlation was reproducibly detected between FOSL and NT5E expression, which suggests their interlinked participation in keeping inflammatory responses in check (Fig. S3E and Fig. S4B). Further, co-depletion of FOSL1 and FOSL2 altered protein-level expression of CD70, STAT4, APOD and JUNB, all of which have reported links to Th17-regulation (*2, 6, 15, 41, 44–46, 70*) (Fig. S3E and Fig. S4A, C, D). These results thus connote a concrete involvement of FOSL proteins in moderating Th17 transcriptional networks.

### FOSL1 and FOSL2 share occupancy over their co-regulated Th17 gene-targets

AP-1 proteins function as transcriptional regulators by directly binding to the target gene loci. To elucidate the global occupancy profiles of FOSL1 and FOSL2 in human Th17 cells, we performed chromatin immunoprecipitation, followed by sequencing (ChIP-seq) analysis. Since these factors portray cell type-specific cellular localization (*71, 72*), immunofluorescence analysis was used to first confirm their predominant nuclear profile in Th17 cells (Fig. 4A). Our ChIP-seq analysis identified 22,127 peaks for FOSL2 and 4,088 peaks for FOSL1 (with irreproducible discovery rate (IDR) significance of < 0.01). In agreement with previous findings, a large fraction of these peaks covered intergenic/intronic regions, thereby suggesting that these factors control gene-expression through distal regulatory elements (Fig. S5A) (*73–75*). Interestingly, comparing the peak distribution profiles of the two proteins revealed a close similarity (Fig. 4B). Further, known FOSL2-binding motifs were detected within FOSL1 peaks and vice-versa, which underscores their propensity to bind to overlapping regions (Fig. 4C). We additionally performed de-novo motif enrichment analysis to identify the consensus DNA-binding sequences of these TFs (Fig. 4C).

**Fig. 4.**
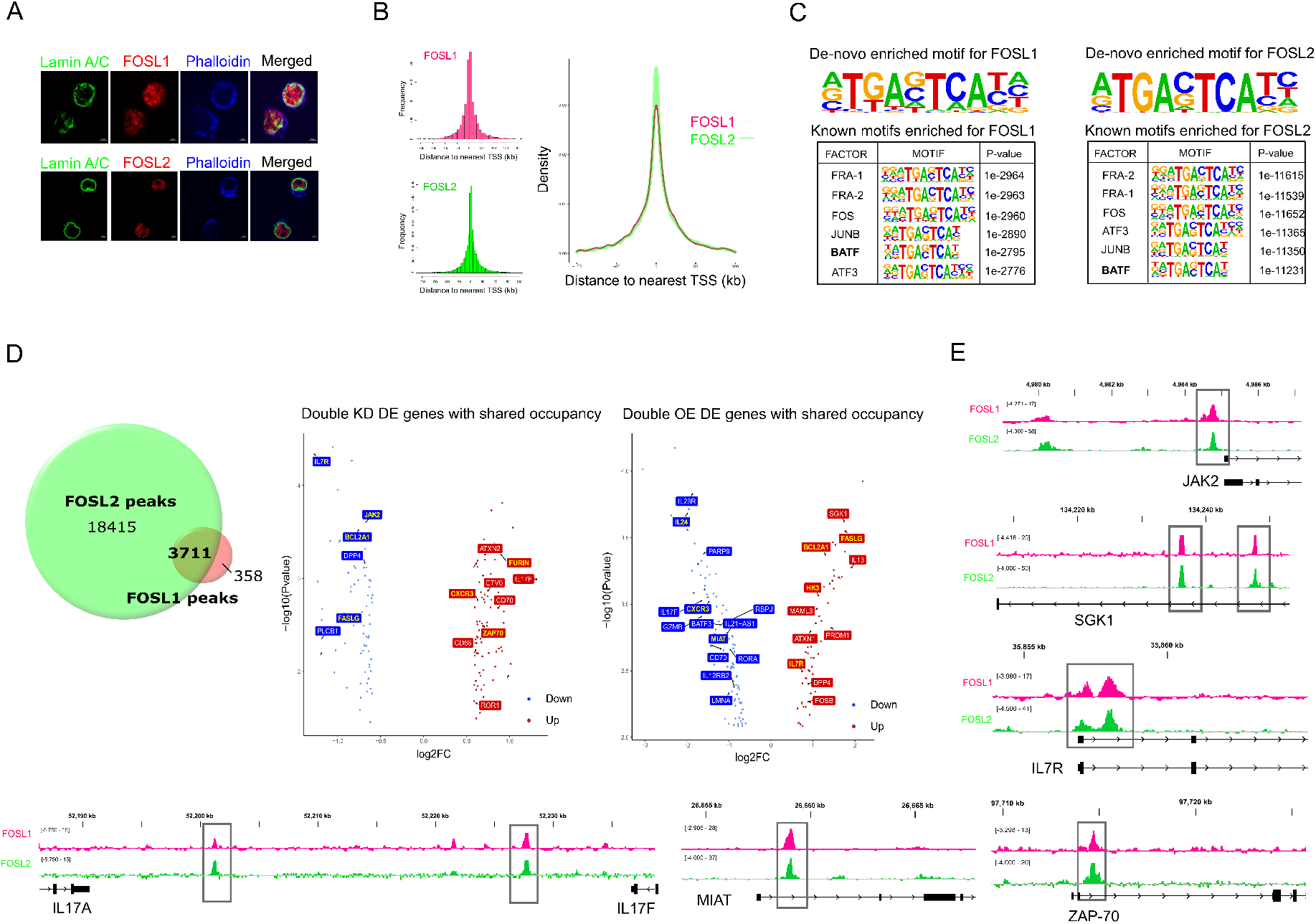
Genome-wide occupancy profile of FOSL proteins in human Th17 cells. (**A**) Immunofluorescence images showing nuclear localization of FOSL1 (red, above panel) and FOSL2 (red, below panel) in Th17 cells polarized for 72h. Lamin A/C (in green) marks the nuclear periphery, whereas phalloidin (in blue) stains the cytoplasmic actin. (**B**) ChIP-seq analysis was performed for FOSL1 and FOSL2 using Th17 cells cultured for 72h. Figures on the left show distribution of FOSL1 and FOSL2 binding sites relative to the position of the closest transcription start site (TSS). TSS is defined to be at position zero. The adjoining figure on the right is an overlay plot that compares the binding profiles of the two factors. (**C**) The topmost consensus sequences for FOSL1 and FOSL2 genomic-binding were identified using *de-novo* motif enrichment analysis by Homer. FOSL1 [left] and FOSL2 [right] peaks were further enriched for known TF motifs and the top six motifs identified by Homer are shown. Peaks with IDR p < 0.01 were used for motif discovery. (**D**) ChIPpeakAnno was used to determine the overlap in the genomic binding sites of FOSL1 and FOSL2 (overlap represents peaks sharing 200 bp or more). Genes neighboring to these overlying sites and differentially expressed under DKD or DOE conditions (FDR ≤ 0.1, |fold-change| ≥ 1.5) were assigned as the shared-direct targets of FOSL1 and FOSL2. Adjoining volcano plots show the logarithmic fold changes for selected shared targets (DKD [left] and DOE [right]). Downregulated genes are in blue, and upregulated ones are in red. Targets with FOSL occupancy over promoter regions (5-kb window around TSS) are highlighted in yellow. (**E**) Integrative Genomics Viewer (IGV) track snapshots show the binding overlap of FOSL1 and FOSL2 over selected Th17 genes.

FOSL1 and FOSL2 are reported to co-occupy selective gene targets in breast cancer cells (*32, 33*). We examined if a similar paradigm exists in human Th17 cells, which potentially mediates the synergistic control of these factors over the lineage. Our genome-wide occupancy analysis revealed 3,711 binding sites to be common in FOSL1 and FOSL2 (Fig. 4D). Strikingly, more than 150 genes proximal to these shared sites were co-perturbed by the two factors in our transcriptome analysis (DKD as well as DOE dataset) (Fig. 4D). These were assigned as the shared-direct targets of FOSL1 and FOSL2, and included numerous Th17-relevant genes that were either activated (*IL13* (*62*), *IL7R (36), JAK2 (76), BCL2A1(63), FASLG (34, 35, 77), PRDM1* (*19*)) or repressed (*IL17F, IL23R, FURIN (78), RBPJ (60), CXCR3 (79–81), MIAT (82), IL24 (53–55), ETV6 (1), ZAP70 (83), RORA).* Integrative genomics viewer (IGV) tracks in Fig. 4E illustrate the binding overlap of FOSL factors over a few of their Th17-gene targets.

An intriguing candidate among the shared targets was *PRDM1 (*or *BLIMP1)*, an inhibitor of Th17-differentiation (*19*), that was directly-bound and positively regulated by FOSL1 and FOSL2. We found this to corroborate with findings reported in other cell types (*11, 84*). Such coordinated expression could be executed through two AP-1 binding sites that exist in proximity of the *BLIMP1* promoter (*85*), which potentially facilitate the co-binding of FOSL proteins. We additionally observed that only one-third of the shared targets showed FOSL occupancy over putative-promoter regions (5-kb region around the TSS) (Fig. S5B; Fig. 4D). The remaining majority, were bound over intronic or intergenic sites. This highlights the fact that FOSL1 and FOSL2 co-regulate Th17 gene-expression programs, presumably by occupying enhancer or silencer elements in the genome.

### FOSL proteins and BATF co-localize over key Th17 genes and regulate their expression in an opposite fashion

Genomic co-occupancy is a distinguished feature of FOS, JUN and ATF family members (*1, 2, 86*). In light of this, the FOSL1 and FOSL2 ChIP peaks were screened for the presence of other known TF-motifs. Our analysis revealed binding motifs for BATF, JUNB, FOS and ATF3, among the top identifications (Fig. 4C). A former study suggested an antagonistic relationship between BATF and FOSL2, during murine Th17-differentiation (*1*). We aimed at verifying whether BATF similarly interplays with FOSL1 and FOSL2, while regulating human Th17 responses.

BATF is a key-modulator of murine Th17 fate (*5, 87*), however its role in the human counterpart remains unknown. This was addressed using RNAi, where naive CD4^+^ T cells were nucleofected with BATF-targeting siRNA and further polarized to Th17 phenotype. Loss of BATF significantly reduced CCR6 and IL-17 levels at 72h of differentiation (Fig. 5A & B; Fig. S6B & C). Further bolstering these results, transcriptome analysis of BATF KD cells showed downregulated expression of multiple Th17-marker genes such as *IL17A, IL17F, IL23R, CCR6* and *IL21* (Fig. 5C and S6A). Additionally, Ingenuity Pathway Analysis found BATF to alter genes involved in IL-23 signaling, T-helper cell differentiation, Th17 activation, and autoimmune processes (SLE, RA) (Fig. 5D). We then examined the global occupancy profile of BATF by performing ChIP-seq analysis on Th17 cells cultured for 72h. A total of 16,479 binding sites (IDR significance < 0.01) (Fig. 5E) were identified, which included putative promoters of more than 4000 genes. Out of these, 35 genes appeared to be transcriptionally regulated by BATF, including multiple lineage-associated targets (*IL21, CCR6, PRDM1, IL17A* and *FASLG*) (Fig. 5G). Adjoining IGV images in Fig. 5G illustrate the occupancy of BATF over promoters of key Th17 genes. Further, motif analysis of the ChIP-seq peaks revealed BATF as the topmost known-motif and also identified the consensus sequence for its genomic-binding (Fig. 5F)

**Fig. 5.**
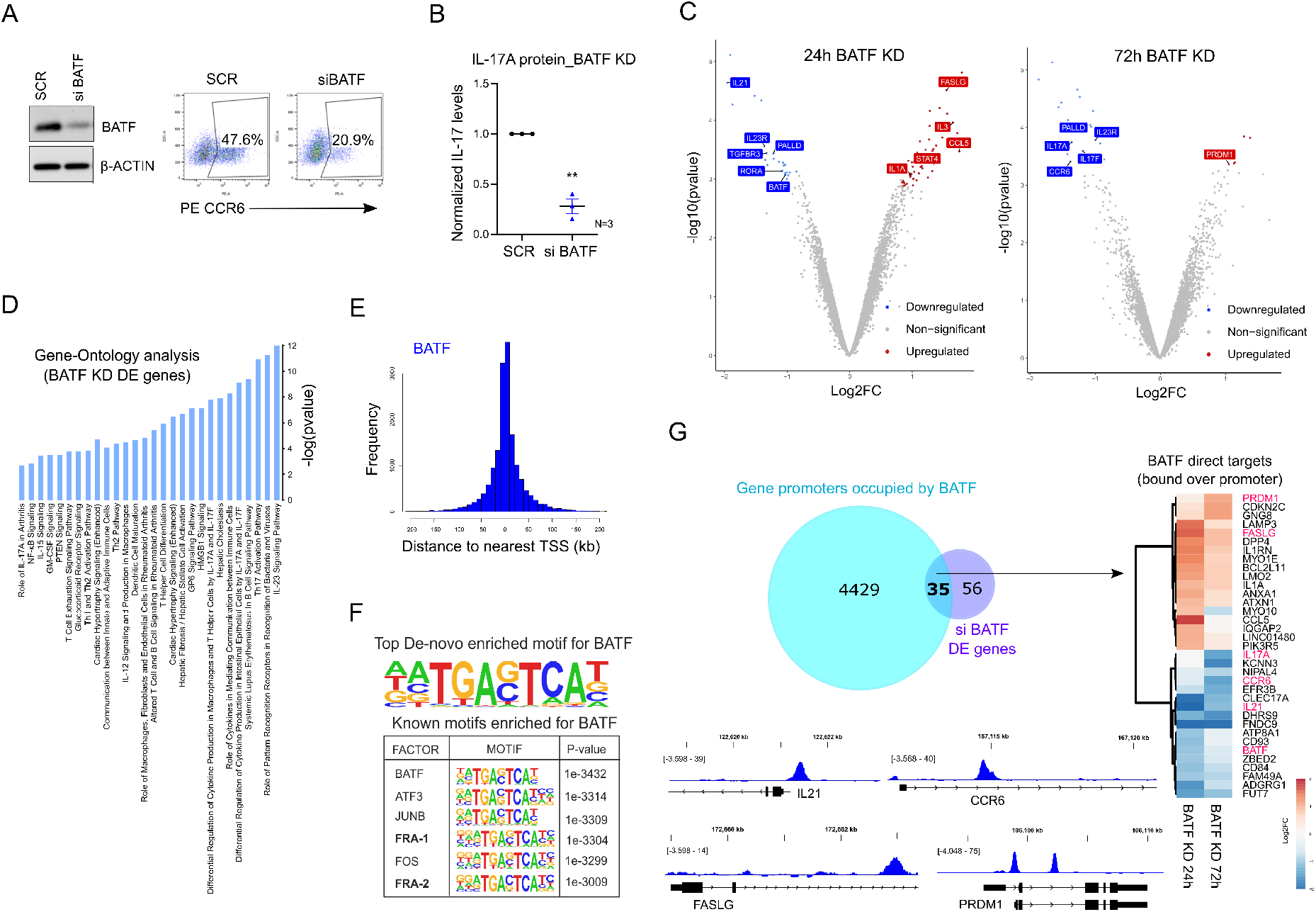
Loss of BATF impairs Th17 differentiation. (**A** and **B**) Immunoblot shows BATF protein levels in control (SCR) versus BATF KD Th17 cells at 24h of differentiation. Adjoining flow cytometry plots depict the percentage of CCR6 positive cells in the indicated conditions, at 72h of Th17 polarization. Panel B shows ELISA analysis for secreted IL-17A levels in SCR versus BATF KD Th17 cells (72h). ELISA values were first normalized to live cell count, followed by normalization with SCR. Graph shows mean ± SEM for three biological replicates. Statistical significance was calculated using two-tailed Student’s t test (**p < 0.01). (**C**) Volcano plots highlight the significantly upregulated (red) and downregulated (blue) genes in BATF-silenced Th17 cells at 24h [left] and 72h [right] of polarization (FDR ≤ 0.1, |fold change| ≥ 1.8). DE genes with relevance to Th17 function are shown. (**D**) IPA was used to identify pathways altered upon silencing of BATF in Th17-polarized cells. (**E**) ChIP-seq analysis was performed for BATF using Th17 cells cultured for 72h. Figure shows distribution of BATF binding sites relative to the position of the closest transcription start site (TSS). TSS is defined to be at position zero. (**F**) The topmost consensus sequence for genomic-binding of BATF and the top six TF motifs enriched within BATF-bound sites, were obtained using de-novo motif enrichment analysis by Homer. Peaks with IDR p value < 0.01 were used for motif discovery. (**G**) Venn diagram shows the overlap between the genes that are altered upon BATF KD and the genes whose promoters are bound by BATF (5-kb around TSS). The overlapping area represents the promoter-bound regulatory targets of BATF and the adjoining heatmap shows their corresponding logarithmic fold change values in BATF KD Th17 cells. IGV images illustrate the occupancy of BATF over some of its Th17-associated targets.

The above findings collectively establish that BATF positively regulates early Th17 differentiation in human, and thus exhibits functions antagonistic to FOSL proteins. To dissect this antagonism at the level of gene targets, we compared the DE genes for BATF KD and FOSL DKD, and focused on the candidates that were common but regulated in an opposite fashion (Fig. 6A, top panel). Likewise, the genes that were similarly altered between BATF KD and FOSL DOE were selected (Fig. 6A, bottom panel). Based on our analysis, the Th17-promoting genes that were negatively regulated by FOSL proteins (*IL17A, IL17F, IL21, RORA, IL23R and CCR6),* were found to be positively regulated by BATF. Concurrently, the Th17-repressor genes that were activated by FOSL factors (*PRDM1 (19) and ID3 (64))*, appeared to be supressed by BATF.

**Fig. 6.**
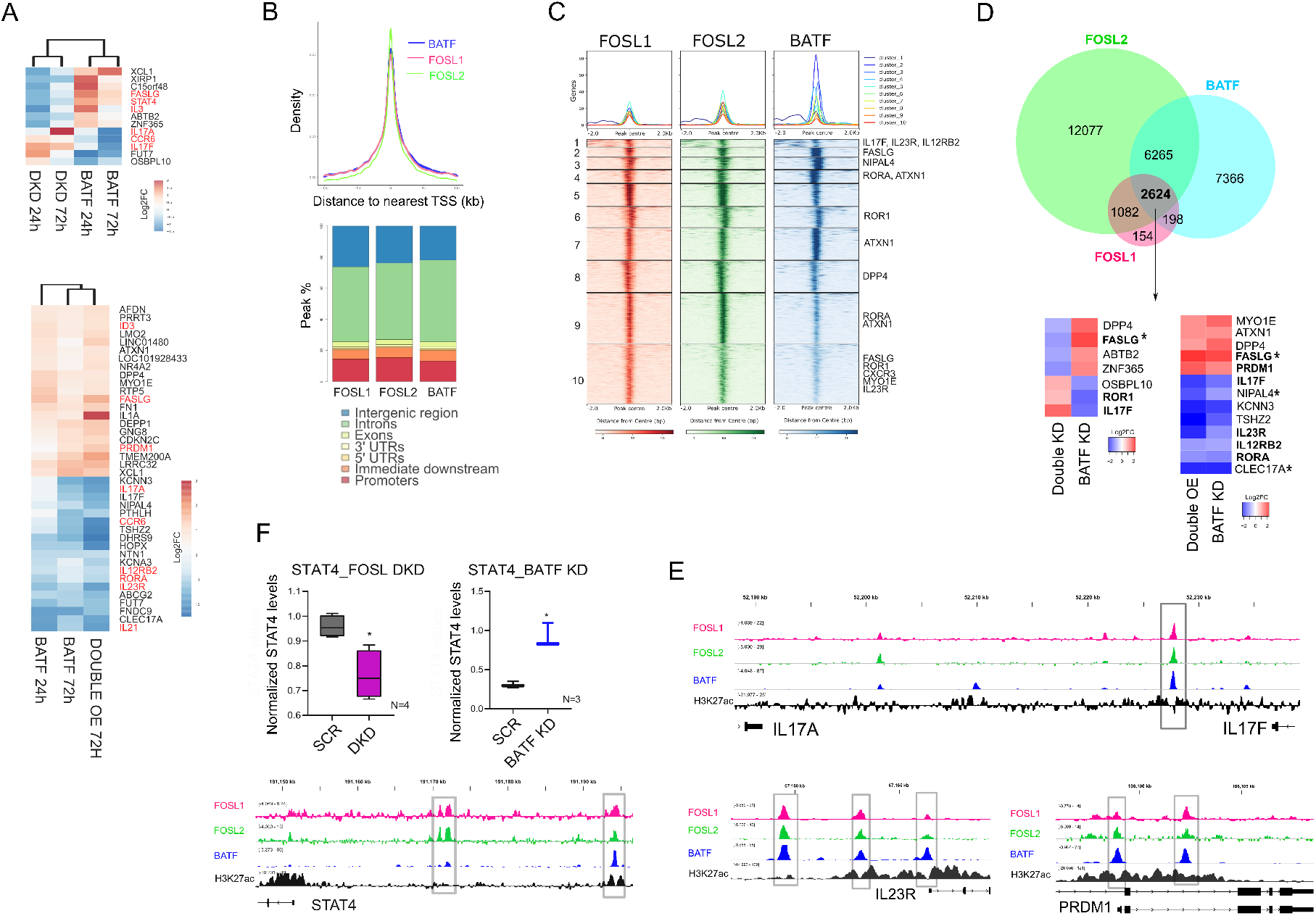
Comparing transcriptional targets and genomic binding sites of FOSL proteins with BATF. (**A**) Heatmap on the top shows logarithmic fold-change values for the DE genes that are oppositely regulated in FOSL DKD and BATF KD Th17 cells, at the indicated time points of polarization. Heatmap in the bottom panel depicts the DE genes that are similarly altered in FOSL DOE and BATF KD Th17 cells. Th17-relevant genes are highlighted in red. (**B**) Comparing the ChIP-seq profiles of FOSL1, FOSL2 and BATF in Th17 cells. Graph (above) shows the overlay between the peak distribution profiles of the three TFs. Bar plot (below) depicts peak-annotation results for their identified binding sites. (**C**) Heatmap with k-means clustering shows the ChIP-seq signal intensities ± 2-kb around the centers of the genomic-binding regions of FOSL1, FOSL2 and BATF. Th17-associated genes in the vicinity of the binding sites are highlighted within the respective clusters. (**D**) Venn diagram shows an overlap between the genomic binding sites of FOSL1, FOSL2 and BATF (overlap represents peaks sharing 200 bp or more). Adjoining heatmap depicts fold-change values for the gene targets that are co-bound and oppositely regulated by FOSL proteins and BATF. Genes showing shared occupancy of the three factors over putative-promoters have been marked (*asterisk). Th17-relevant targets are highlighted. (**E**) IGV track snapshots illustrate the co-localization of FOSL1, FOSL2 and BATF over selected Th17 genes. Profile of H3K27ac marks around the shared sites is shown. (**F**) Bar plot depicts immunoblot-based expression analysis of STAT4 in FOSL DKD [left] and BATF KD [right] Th17 cells, cultured for 72h. Data shows mean ± SEM for three or four biological replicates, as indicated. Statistical significance is calculated using two-tailed Student’s t test (*p < 0.05). Adjoining IGV track shows the binding overlap of FOSL1, FOSL2 and BATF, flanked by H3K27ac marks near the STAT4 locus.

To further examine which of the genes regulated in an opposite fashion are also bound by these TFs, we compared the ChIP-seq profiles of the three proteins. An overlay of their peak distribution plots suggested their co-localized occupancy (Fig. 6B, top). In addition, mapping of the ChIP-seq signal intensities ± 2-kb around the centres of the genomic-binding regions of FOSL1, FOSL2 and BATF, evidently demonstrated the unanimity in their DNA binding pattern (Fig. 6C). Their individual sites were further clustered; where clusters 2-4 and 7 showed higher enrichment of BATF, whereas clusters 5, 8, 9 & 10 depicted greater signal densities for FOSL1 and FOSL2. We then used the R package ChIPpeakAnno (*88*) to determine the exact overlap in their binding sites and found a total of 2,624 sites to be common (Fig. 6D). Interestingly, several genes neighbouring to these common binding sites were antagonistically regulated by FOSL and BATF (Fig. 6D). Multiple genes controlling Th17 effector-functions (*RORA, IL17F, ROR1, IL23R, PRDM1, FASLG, IL12RB2*) were included within this group (highlighted in Fig. 6D). We thus propose that BATF and FOSL contextually or competitively bind to a common set of Th17-defining genes, and oppositely regulate lineage-establishment. Moreover, the concerned factors primarily occupy regulatory DNA elements distal to the promoter in order to influence their targets (Fig. 6B (bottom) and Fig. 6D (excluding the (*) genes)).

Many AP-1 TFs show co-localized binding on genomic regions that have enhancer marks (H3K4me1/H3K27ac). Such regulatory modules are known to drive cellular differentiation and disease-associated functions (*74, 89*). To scrutinize our study on this front, we examined the ChIP peaks of FOSL1, FOSL2 and BATF for H3K27Ac marks (a transcriptionally permissive histone modification found on active enhancers and promoters), by using a published human Th17 dataset ((*90*); GSE101389). IGV tracks in Fig. 6E illustrate how H3K27ac flanks the shared binding sites of these AP-1 factors, in the vicinity of their direct targets *(IL17A/F, IL23R* and *PRDM1*). An identical pattern was observed upstream of the human *STAT4* locus (*IGV image*, Fig. 6F). Despite its well-established role in Th1 polarization (*42, 91*) and autoimmunity (*92*), STAT4 has not been studied in the context of non-pathogenic human Th17 responses. To validate the effects of FOSL and BATF on STAT4 expression, we performed immunoblot analysis. Our results revealed that loss of BATF upregulated STAT4 levels, whereas co-depletion of FOSL1 and FOSL2 reduced STAT4 expression (Fig. 6F; Fig. S6D). The concerned AP-1 factors may thus instrument Th17 responses by fine-tuning the lineage-diversification to Th1 fate.

FOS, JUN and ATF proteins can bind to common interacting partners. This possibly creates molecular competition, which is known to mediate functional antagonism between specific members of the AP-1 family (*21, 93–96*). BATF for instance, competes with FOS proteins for partnering with JUNB, which allows it to negatively influence FOS activity (*94*). To address if a similar mechanism facilitates the BATF-FOSL antagonism in our study, it was important to primarily check for their common binding partners. The interactomes of FOSL1 and FOSL2 in human Th17 cells were recently uncovered by a parallel study from our lab, using global proteomics approaches (*97*). The two proteins were found to share numerous binding partners, many of which regulate T cell signaling processes (RUNX1, SIRT1, EIF4E, JUN, JUNB, ADAR, NUFIP2, HSPH1, IFI16, HNRNPH1/2, LARP4 and DHX9) (Fig. S7A). Out of these, JUN TFs are already reported to interact with BATF, in other studies (Fig. S7B). To verify these findings in human Th17 cells, we immunoprecipitated BATF from 72h-polarized populations and performed immunoblot analysis (Fig. S7C). In addition to JUN (JUN, JUNB), BATF interaction was also checked for other shared partners of FOSL1 and FOSL2 (RUNX1 and SIRT-1). Our results found BATF to reliably associate with JUN, JUNB and RUNX1 (Fig. S7C), all of which are established regulators of Th17-differentiation (*2, 3, 6, 98, 99*). Interestingly, these identified interactors are known to exhibit context-specific roles, based on their choice of binding partner (*21, 99*). This implies that BATF and FOSL factors potentially compete to interact with common proteins, and differentially orchestrate human Th17 responses. Notably, STAT3 and IRF4, which form pioneering complexes with BATF in mouse Th17 cells (*1, 100*), showed no interaction with it in the human counterpart (Fig. S7D). This may hint at subtle differences in the early signaling events of the two species.

### Multiple disease-linked SNPs were enriched within the genomic binding sites of FOSL1, FOSL2 and BATF

Functional analysis of data from genome-wide association studies (GWAS) has revealed that SNPs linked to disease phenotypes can alter binding sites of key TFs (*101*). The presence of a SNP can abrogate or enhance TF occupancy, which might subsequently influence gene-expression profiles (*26*). Interestingly, 90% of the disease-linked SNPs are reported to occur within non-coding genomic regions (*102*), which also appear to accommodate a major fraction of the TF ChIP peaks in our study. With this in view, we sought to determine whether the genomic-binding sites of FOSL1, FOSL2 and BATF harbour any autoimmune-associated SNPs that could disrupt the occupancy of the respective factors.

We used the NHGRI-EBI GWAS catalogue from Caucasian populations to primarily query SNPs with reported links to 11 different autoimmune phenotypes (celiac disease, IgA immunodeficiency, RA, ankylosing spondylitis (AS), Crohn’s disease (CD), MS, psoriasis (PS), primary biliary cholangitis, type I diabetes and ulcerative colitis (UC) (Fig. 7A). Upon intersecting these with the TF peaks identified in our study, we detected 114, 571 and 573 disease-linked SNPs (and their proxies) within FOSL1, FOSL2 and BATF binding sites, respectively. Importantly, the genomic binding regions shared between the three factors harboured as many as 64 disease-associated SNPs.

**Fig. 7.**
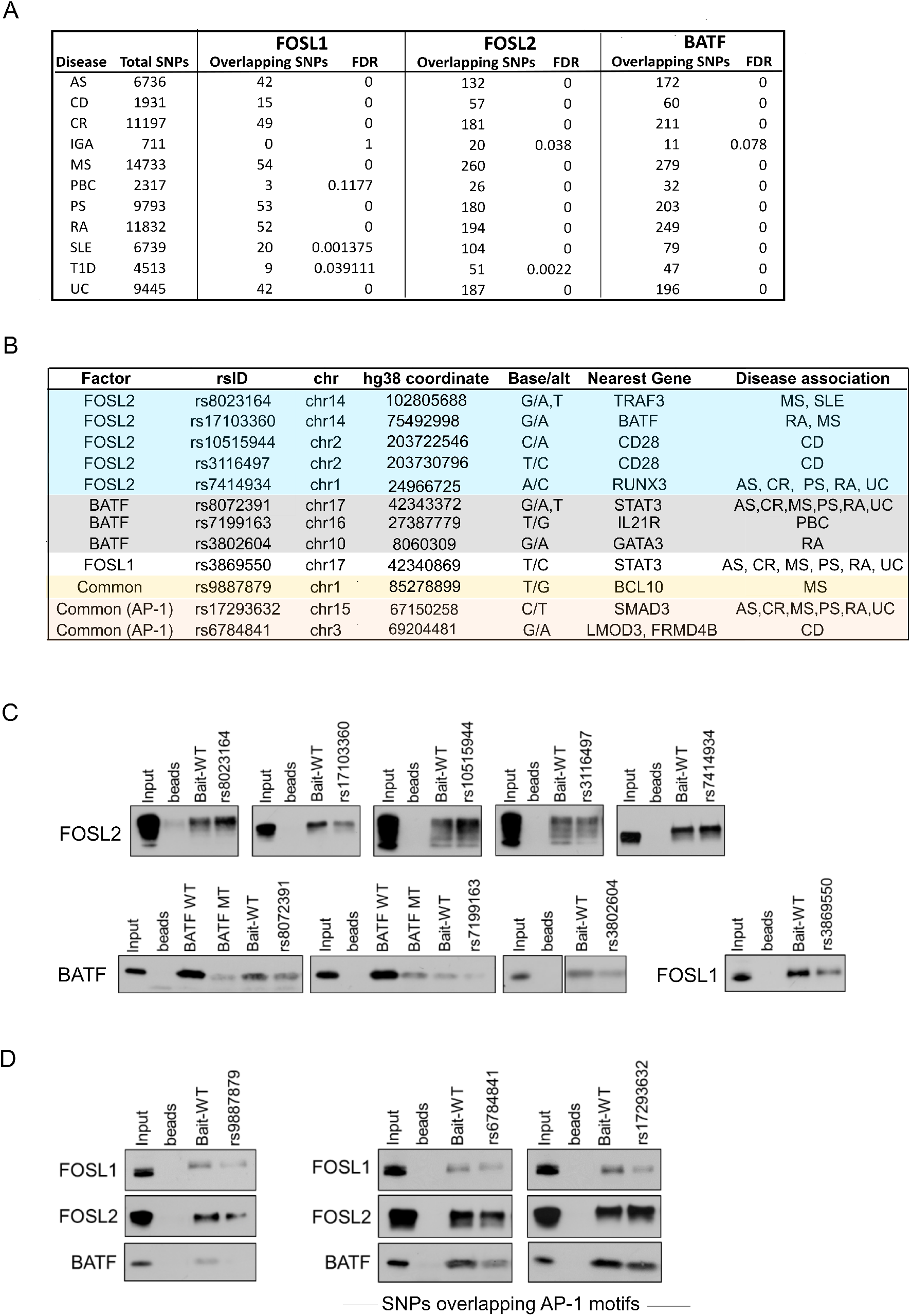
Autoimmune SNPs within the genomic binding sites of FOSL1, FOSL2 and BATF influence the ability of these factors to bind DNA. (**A**) Enrichment of disease-associated SNPs (or their proxies in Caucasian populations) within FOSL1, FOSL2 and BATF genomic-binding sites, relative to random sets of background SNPs. (**B**) SNPs relevant to the study were shortlisted. Out of these, the SNPs that were functionally validated in DNA-affinity precipitation assays (DAPA) have been shown. (**C** and **D**) DAPA reveals the SNPs that alter the binding of FOSL1, FOSL2 or BATF to their genomic sites that were identified by ChIP-seq analysis. Wildtype (WT) oligonucleotides containing the binding motifs of these TFs (at different genomic loci), and mutant oligonucleotides harboring a SNP within the corresponding motif, were used as baits. For experimental controls, an oligonucleotide with a conserved binding sequence for BATF (BATF WT), and the corresponding mutated sequence which is known to disrupt BATF occupancy (BATF MUT) were used. Immunoblot results for the SNPs unique to FOSL1, FOSL2 and BATF (panel C), and the ones common across the three factors (panel D) are shown. Data is representative of three biological replicates.

We further shortlisted the SNPs relevant to our study by screening for the ones that overlap with the TF binding sites in the vicinity of Th17-relevant genes. Additionally, the SNPs that were common across the three factors and harboured within canonical AP-1 motifs were listed (Fig. S8A). DNA-affinity precipitation assay was then performed to determine if any of these SNPs affect the binding of FOSL1, FOSL2 or BATF to DNA. Wildtype oligonucleotides with TF binding motifs (identified from FOSL1, FOSL2 or BATF ChIP-seq) and mutant oligonucleotides with the SNP introduced within the motif, were used as baits for the assay. DAPA analysis of selected SNPs (Fig. 7B) revealed important changes in the binding affinities of the three TFs. For instance, we detected altered binding of FOSL2 to mutant oligonucleotides harbouring the following five SNPs: rs8023164 (MS and SLE), rs17103360 (MS and RA), rs10515944 (CD), rs3116497 (CD) and rs7414934 (AS, CR, PS, RA, and UC) (Fig. 7C; Fig. S8B). These SNPs appeared to occur in the regulatory regions that are neighbouring to *TRAF3, BATF, CD28* and *RUNX3 genes,* which could be potential targets of FOSL2. Interestingly, TRAF3 is reported to enhance T cell activation (*103*), restrain IL-2 dependent generation of thymic Tregs (*104*) and impair IL-17R proximal signaling (*105*). BATF is a well-known regulator of Th17 responses (*5*), whereas low CD28 co-stimulation has been found to promote Th17-development (*106, 107*). Furthermore, while RUNX transcription factors are recognized modulators of Th17-fate (*108*), RUNX3 in particular, is found to be elevated in CD4^+^ T cells of PS patients. Notably, loss of RUNX3 impairs Th17 and Th22 differentiation, both of which are required for the pathogenesis of psoriasis (*109*). Our findings thus suggest that specific SNP mutations alter the ability of FOSL2 to bind to target regulatory DNA elements near important Th17-signaling genes.

We similarly identified three SNPs for BATF (near *IL21R, GATA3* and *STAT3*) and one for FOSL1 (near *STAT3*), which when introduced within the corresponding TF motif, significantly disrupted occupancy of these factors (Fig. 7C; Fig. S8B). IL-21 and STAT3 positively regulate Th17 cell programs (*26, 49, 57, 110, 111*), whereas GATA-3 is a master regulator of Th2-fate, which also constrains Th17-mediated pathology (*112, 113*). Interestingly, a BCL10-proximal SNP rs9887879, which overlaps the shared binding sites of FOSL1, FOSL2 and BATF, reduced the DNA-binding affinities for all of them (Fig. 7D; Fig. S8C). BCL10 suggestively regulates Th17 function as a part of a signaling complex (*114*). It is a key component of the Carma1-Bcl10-Malt1 complex that is essential for pathogenic Th17 responses (*115*). We additionally validated the functional effects of two other SNPs – rs17293632 near *SMAD3* (linked to AS, CR, MS, PS, RA and UC) and rs6784841 near *LMOD3/FRMD4B* genes (linked to CD) (Fig. 7D; Fig. S8C). These occurred within consensus AP-1 motifs at the shared genomic-binding regions of the three factors, and significantly altered the binding propensities for all of them. The ability of the above-mentioned SNPs to perturb genomic occupancy of these TFs could trigger changes in their Th17-regulatory roles, thereby facilitating the development of multiple autoimmune phenotypes.

## DISCUSSION

FOS and ATF proteins are established regulators of proliferation, differentiation and apoptosis in many cancers. Their involvement in lineage specification of T-helper cell types, however, has been investigated only recently. Th17-specific AP-1 networks have been mostly studied in mouse models. Taking into account the recently established heterogeneity between human and mouse Th17 cells (*15*), we used human T cells to verify the roles of FOSL1 and FOSL2 during early stages of Th17 differentiation. Using complementary approaches, we demonstrated that both factors negatively influence Th17-induction, thereby highlighting that FOSL1 has different roles in human and mouse (*3*).

AP-1 factors co-ordinate with each other to drive gene-expression programs (*21*). FOS proteins, in particular, exhibit functional redundancy that allows them to compensate for the loss of each other (*21, 116, 117*). Data from the present study revealed that individual perturbation of FOSL1 or FOSL2 only modestly altered Th17 cell-identity. Disrupting them in parallel, however, caused additive changes in gene-expression. Remarkably, co-depletion or dual over-expression of the two proteins cooperatively affected several genes associated with Th17 cell-function (*IL17A, IL17F, NT5E* (*37, 38, 40*)*, CCR6, IL7R (36), IRF7 (56), BCL2A1 (63), DUSP2 (48), PRDM1* (*19*)*, IL21* (*57*)*, JUNB* (*2, 6*)*, IL23R, CXCR3* (*79*)*, IL12RB1 (49), CD52 (58, 59), TIGIT* (*65*)*, ID3* (*64*)). Our findings thus confirm that these paralogs jointly instruct the initial stages of human Th17 differentiation.

Previous studies in mouse indicate that FOSL2 suppresses Th17-responses, yet promotes the expression of genes involved in sustenance of the lineage (*1*). Our results, however, portray a different scenario. Genes associated with Th17-maintenance (*Il23r, Il12rb1 and Il21*) that were activated by FOSL2 in mouse (*1*), were in fact inhibited by it in the human counterpart. This implies that although FOSL2 similarly represses Th17 cell-effector genes in the two species, its involvement in parallel signaling networks may differ in human and mouse. FOSL proteins were additionally found to co-influence multiple genes involved in the development of other T-helper cell fates, including *TBX21, GATA3, IFNG, FURIN, BATF3, IL12RB2, HOPX and IL13*. For instance, FOSL1 and FOSL2 negatively-regulated Th1-lineage genes (*TBX21 (118, 119), IFNG (120), BCL3 (121), IL12RB2 (122), HOPX (123)*), while promoting the expression of Th2-specific factors (*GATA3 (112), IL13 (124))*. It however remains to be determined whether FOSL proteins truly restrain Th17 responses by modulating helper-T-cell diversification.

Though murine studies have examined the molecular networks that drive the transition from homeostatic-to pathogenic-Th17 fate, this switch is poorly understood in human. Our transcriptome analysis revealed FOSL1 and FOSL2 to jointly suppress several genes that positively correlate with Th17-pathogenicity, including *GZMB (125), IL23R (126, 127), RBPJ (60), IFN-γ (128, 129), and TBX21 (108, 125, 130)*. In addition, FOSL factors downregulated *IL-26* expression, a cytokine that marks inflammatory Th17-populations in patients suffering from Crohn’s disease (*131*). They also inhibited the expression of *FGF2*, which coordinates with IL-17A to drive autoimmune arthritis (*47*). These findings suggest that FOSL proteins could help in retaining the protective nature of Th17 cells, under conditions of adversity. Furthermore, they affected the expression of several receptors/ligands including *CCL3L3 (132), CCL4 (133), CXCL8 (134), CXCR3 (79–81), and CCR6 (68)*, that govern the migration of inflammatory T cells in autoimmune phenotypes. Further investigation on how FOSL1 and FOSL2 modulate pathogenic Th17-signaling, could define their potential in the treatment of relevant diseases.

STAT3 acts as a master-regulator of Th17-differentiation in both human and mouse (*26, 44, 135–138*). We found FOSL proteins to inhibit several genes that are known to be activated by STAT3 (*HOPX, IL23R, CCR6, IL24, FNDC9, GZMB)* (*26*). Despite their opposite roles in controlling Th17-fate, STAT3 positively regulated FOSL expression in human Th17 cells. The existence of a STAT3-based mechanism to induce these Th17-inhibitors could be explained through a recent study, where STAT3 was found to alternatively drive a negative-feedback loop that limits Th17-mediated tissue damage (*139*).

The functional antagonism between FOSL2 and BATF is well-reported in mouse (*1*). Our study is the first one to investigate the relationship between these factors in human Th17 cells. Our findings further reveal how BATF function diverges from FOSL1 at the level of transcriptional regulation, which has not been addressed before. BATF and FOSL factors were found to directly bind and oppositely-regulate key Th17 marker genes (*IL17A, IL17F, IL23R, CCR6, IL21*), along with other candidates that are associated with the lineage (*IL3, STAT4, FASLG, PRDM1, IL12RB2* and *RORA)*. A cardinal target among these was *FASLG,* which is a crucial regulator of apoptosis *(34, 35, 77)*. We found its expression to be driven by FOSL proteins and inhibited by BATF. Responsiveness to FAS-signaling contextually varies for pro-inflammatory and anti-inflammatory cells, and is reported to subsequently decide whether autoimmunity develops (*34, 140, 141*). Insights on AP-1-governed FAS networks could thus hold significance in disease-biology.

Our analysis found BATF to regulate many Th17-lineage genes by occupying their promoter-regions. However, the BATF-bound sites that co-localized with FOSL near their oppositely-regulated targets, mostly occurred within intergenic or intronic elements. In mouse Th17 cells, several AP-1 factors are known to co-bind their consensus motifs within the intergenic regions of *Il17a/f* loci (*2*). This paradigm appears to be conserved, since we discovered FOSL1, FOSL2 and BATF to similarly intersect over the corresponding loci in human Th17 cells. Many studies including ours, indicate that such binding convergence occurs over enhancer landscapes (*74, 86, 142*), which potentially govern lineage-identity and plasticity of T-helper cell fates (*143, 144*). Nonetheless, it remains to be understood whether polarization-induced epigenomic changes are guided by these AP-1 factors in human T cells.

The shared genomic occupancy of FOSL and BATF warrants further investigation. Although the overlap in their ChIP peaks suggests co-occupancy or competitive binding, additional experiments are required to ascertain the precise mode of their action. For instance, competitive-binding of these TFs could be confirmed through gene-perturbation approaches, provided that either of these factors show an enhanced occupancy in absence of the other. Such findings have previously been reported for JUNB and JUND, which also exhibit functional antagonism in Th17 cells (*2*). Although our results highlight the BATF-FOSL interplay in regulation of Th17-effector functions, follow-up studies are required to address their cross-talk in early signaling events. BATF acts as a pioneer factor that mediates nucleosomal clearance at lineage-associated loci, during the induction of T cell differentiation. The role of FOSL proteins in this context, however, is poorly investigated (*145*). Studying the temporal dynamics of their functions is a pre-requisite to understanding the interlinked roles of these AP-1 factors.

We identified hundreds of autoimmune disease-associated SNPs within the genomic-binding sites of FOSL1, FOSL2 and BATF. An ongoing study from our lab further revealed that a large fraction (60–80%) of these binding sites overlap with Th17-specific enhancers. Remarkably, the enhancer-specific binding regions of FOSL1, FOSL2 and BATF harboured as many as 100, 470 and 478 disease-linked SNPs, respectively (unpublished data). Notably, over 50 of these SNPs were common across the three factors. These single-base changes could alter AP-1 function at distal regulatory elements that dictate Th17 gene expression programs. Furthermore, selected SNPs relevant to our study altered the binding abilities of FOSL1, FOSL2 and BATF in *in vitro* DNA-binding assays. Regardless, these effects need to be confirmed by generating SNP knock-in clones and examining their *in vivo* influence on the genomic occupancy of these factors, as compared to the reference clones. This could help in dissecting the subsequent changes in FOS and ATF-mediated transcriptional regulation of Th17 cells, which potentially contributes to the development of autoimmunity.

## MATERIALS AND METHODS

### Study design

The objective of this study was to advance our understanding on the transcriptional mechanisms that regulate human Th17-differentiation. Naive CD4^+^ T cells isolated from human umbilical cord blood of neonates were *in vitro* differentiated to obtain Th17 cells, which was the primary cell source used in this study. Transient perturbation of gene-function was achieved using RNAi silencing as well as RNA-based over expression strategies. Global transcriptional targets and genome-wide binding sites of the candidate TFs were determined using RNA-seq and ChIP-seq analysis. Flow-cytometry, immunoblotting, qRT-PCR and ELISA methods were used to assess the differentiation of naive CD4^+^ T cells to Th17 phenotype, as well as to validate the observations from the sequencing datasets. Autoimmune-disease linked SNPs were queried within TF-binding sites using GWAS catalogue and the functional effects of selected SNPs were determined using DNA-affinity precipitation assays. Information on experimental replicates have been provided in the adjoining legends of each figure panel. Further details on methods and analysis have been provided in the Supplementary information. This research on primary human CD4^+^ T cells from the cord blood of neonates was conducted only after approval from the joint ethical committee of University of Turku and Turku University Hospital.

### Primary human CD4^+^ T-cell isolation and Th17 culture

Human cord blood mononuclear cells (CBMCs) were isolated from the umbilical cord blood of healthy neonates (Turku University Central Hospital, Turku, Finland) using the Ficoll-Paque density gradient centrifugation (Ficoll-Paque PLUS; GE Healthcare). Naive CD4^+^ T cells were further purified using CD4^+^ Dynal positive selection beads (Dynal CD4 Positive Isolation Kit; Invitrogen). CD4^+^ T-cells were stimulated with plate-bound α-CD3 (3.75 μg/ml; Immunotech) and soluble α-CD28 (1 μg/mL; Immunotech) in X-vivo 20 serum-free medium (Lonza). X-vivo 20 medium was supplemented with L-glutamine (2 mM, Sigma-Aldrich) and antibiotics (50 U/mL penicillin and 50 μg/mL streptomycin; Sigma-Aldrich). Th17 cell polarization was induced using a cytokine cocktail of IL-6 (20 ng/mL; Roche), IL-1β (10 ng/mL) and TGF-β (10 ng/mL) in the presence of neutralizing anti-IFN-γ (1 μg/mL) and anti-IL-4 (1 μg/mL) to block Th1 and Th2 differentiation, respectively. For the control cells (Th0), CD4^+^ T-cells were TCR stimulated with α-CD3 and α-CD28 in the presence of neutralizing antibodies (without differentiating cytokines). All cytokines and neutralizing antibodies used in the study were purchased from R&D Systems unless otherwise stated. All cultures were maintained at 37°C in a humidified atmosphere of 5% (v/v) CO_2_/air.

### RNAi silencing

CD4^+^ T cells from umbilical cord blood were suspended in Opti-MEM I (Invitrogen) and transfected with the respective targeting siRNA using the nucleofection technique of Lonza. Control cells were treated with non-targeting siRNA (Sigma).

#### I. STAT3 and BATF knockdown (KD)

Four million cells were transfected with 6 μg of STAT3- or BATF-targeting siRNA, after which the cells were rested at 37° C for 36–40 h in RPMI 1640 medium (Sigma-Aldrich) supplemented with pen/strep, L-glutamine (2 mM) and 10% FCS. Cells were subsequently activated and cultured under Th17 conditions. For identification of BATF target genes, cells were harvested at 24h and 72h post-induction of polarization. Three biological replicates were prepared, each time, including BATF targeting siRNA and non-targeting control siRNA. Total RNA was isolated and samples were prepared. A pool of two siRNAs was used for silencing BATF.

#### II. FOSL knockdown (KD) and double KD (DKD)

For single KD, four million cells were transfected with 5 μg of FOSL1 or FOSL2-targeting siRNAs, and the rest of the protocol was followed as described in (I). Two different siRNAs were used to target FOSL1 and FOSL2 each. For double KD (DKD), four million cells were nucleofected with 10 μg of FOSL-targeting siRNA (5 μg FOSL1 + 5 μg of FOSL2 siRNA) or 10 μg of Scramble siRNA. FOSL1 or FOSL2 single knockdown (KD) nucleofections were performed for comparison (5 μg FOSL1 or FOSL2 siRNA + 5 μg of control siRNA). The rest of the protocol was followed as described in (I). For identification of global targets – SCR, KD and DKD Th17 cells were harvested at 24h and 72h of polarization. Three biological replicates were prepared and subjected to further downstream analysis.

### FOSL over-expression (OE) and Double OE using in vitro transcribed RNA

#### I. Generating in-vitro transcribed (IVT) RNA

To generate linearized vectors for the IVT reaction, the T7 promoter containing plasmids: empty pGEM-GFP64A, pCMV6-AC-GFP-FOSL1 (Origene, Cat no. RG202104) and pCMV6-AC-GFP-FOSL2 (Origene, Cat no. RG204146), were *in vitro* digested using the restriction enzymes Spe1 (NEB, Cat no. R0133), Xma1 (NEB, Cat no. R0180) and Ssp1 (NEB, Cat no. R3132), respectively. Digestion was performed for 1 h using Cut Smart Buffer (NEB, Cat no. B7204S). Next, using the generated templates, IVT RNA was produced using Cell Script MessageMAXTM T7 ARCA-Capped Message Transcription Kit (Cell Script, Cat. no. C-MMA60710, Madison, WI), by following manufacturer’s instructions. 10 M lithium chloride (LiCl) was used to precipitate the product (−20°C, O/N), followed by 70% ethanol washes (two washes, each followed by a 10-min centrifuge spin) and resuspension in nuclease-free water. The size of the RNA was confirmed using BioRad Experion or Agilent Bioanalyzer at this step. The RNA was further poly-adenylated using Cell script A-Plus™ Poly(A) Polymerase Tailing Kit (Cell Script, Cat no. C-PAP5104H). LiCl precipitation was repeated, and the final pellet was resuspended in nuclease-free water. RNA concentration was determined using a Nanodrop™ detector (Thermo Scientific) and the IVT RNA was stored at −80°C till further use.

#### II. Nucleofection

For double over-expression (DOE), 4 million cells were nucleofected with either FOSL1+FOSL2 IVT RNA (56 pmoles of FOSL1 + 56 pmoles of FOSL2) or control GFP RNA (112 pmoles). Single over-expression controls (OE) for FOSL1/FOSL2 were also maintained (56 pmoles of FOSL1 or FOSL2 RNA + 56 pmoles of control GFP RNA). We ensured equimolar RNA amounts across the different nucleofection conditions. Cells were rested for 16–20 h post-nucleofection and further cultured under Th17 conditions. For identification of global targets – GFP, single OE and double OE Th17 cells were harvested at 72h of polarization. Three biological replicates were prepared and subjected to sample preparation for RNA-sequencing analysis.

### Gene-expression analysis

#### I. RNA Isolation and RNA-Seq Sample Preparation

RNA was isolated (RNeasy Mini Kit; QIAGEN, Cat no. 74104) and given on-column DNase treatment (RNase-Free DNase Set; QIAGEN) for 15 min. The removal of genomic DNA was ascertained by an additional treatment of the samples with DNase I (Invitrogen, Cat no. 18068-015). After RNA quantification (using NanoDrop™ 2000) and quality control (using BioRad Experion or Agilent Bioanalyzer), libraries for RNA-Seq were prepared for three biological replicates. The high quality of the libraries was confirmed with Advanced Analytical Fragment Analyzer (Advanced Analytical Technologies, Heidelberg, Germany) or with Agilent Bioanalyzer, and the concentrations of the libraries were quantified with Qubit^®^ Fluorometric Quantitation (Life Technologies, ThermoFisher). Sequencing was performed at the Finnish Functional Genomics Centre (FFGC) using HiSeq3000 Next-Generation Sequencing platform.

#### II. Alignment and Differential Expression Analysis

50-bp single-end reversely-stranded sequencing reads were checked for quality using FastQC (v.0.11.14) (*146*) and MultiQC (v.1.5) (*147*). High-quality reads were aligned to the human reference genome (hg38) using R (v.3.6.1) (*148*)/ Bioconductor(v.3.9) (*149*) package Rsubread (v.1.34. (*150*) which was also used for producing the gene-wise read counts based on RefSeq gene annotations. Statistical testing and differential expression analysis was performed using Bioconductor package ROTS (v.1.12.0) (*151*). For each comparison, the expressed genes (CPM expression value > 1) in at least 50% of the replicates in one of the compared sample groups were included in the statistical testing. DE genes were identified with cut-offs of false discovery rate (FDR) ≤ 0.1 and absolute fold-change ≥ 1.8 (unless otherwise specified). The DE gene heatmaps were produced using R package pheatmap (v. 1.0.12).

### ChIP-seq analysis

#### I. Sample preparation

CD4^+^ T cells were cultured under Th17 cell polarizing conditions for 72h. Chromatin was prepared from 40–50 million cells using Diagenode Chromatin shearing optimization kit (Cat no. C01010055) and further subjected to sonication using Bioruptor sonicator (Diagenode) to obtain chromatin fragments of 100–500 bps. Fragmented chromatin was incubated with 10-– 12 μg of FOSL1 (Santacruz Biotechnology, Cat no.sc-28310), FOSL2 (Cell Signaling Tech, Cat no.19967) or BATF (Cell Signaling, Cat no. 8638) antibody and incubated with magnetic beads for crosslinking (Dynal Biotech/Invitrogen, Cat no. 112.04). The crosslinks were further reversed (65°C for 12–16 h, mixer conditions), treated with proteinase K and RNase A and then purified using QIAquick PCR purification kit (QIAGEN, Cat no. 28104). DNA libraries were prepared using two biological replicates of each TF ChIP and sequenced on Miseq Nano (Fasteris Life Sciences, Plan-les-Ouates, Switzerland).

#### II. Analysis

75-bp paired-end reads were obtained, and quality control was performed with FastQC (v. 0.11.4) (https://www.bioinformatics.babraham.ac.uk/projects/fastqc/). The adapter sequences present in the raw reads were trimmed using TrimGalore! (v. 0.4.5) (https://www.bioinformatics.babraham.ac.uk/projects/fastqc/), and the trimmed reads were mapped to the hg38 reference genome using Bowtie2 (v. 2.3.3.1) (*152*). Duplicate reads were marked with Picard tools (v. 2.20.2) (https://broadinstitute.github.io/picard/) MarkDuplicates function and reads with mapping quality < 30 were filtered out using samtools (v. 1.9) (*153*). Sample quality was controlled by calculating cross-correlation scores and the non-redundant fraction with phantompeakqualtools (v. 1.2) (*154, 155*) and preseq (v. 2.0) (*156*), respectively. Peaks were called using MACS2 (v. 2.1.0) (*157*), and reproducible peaks were identified using IDR (*158*) with an FDR cut-off of 0.01. R packages ChIPpeakAnno (v. 3.21.7) (*88*), and EnsDb.Hsapiens.v86 (v. 2.99.0) were used to annotate the peaks and identify peaks common to all three transcription factors with a minimum overlap of 200 bp. In addition to the nearest features, the annotation includes any features that overlap the peaks resulting in more than one row per peak for many of the peaks in the excel files provided as supplementary files. The number of peaks common to the transcription factors reported in the main text and in the Venn diagrams, is the minimum number of overlapping peaks. Enriched transcription factor binding site motifs within the peaks were identified by Homer (v. 4.11) using both *de novo* and known motifs. A 200-bp region size was used for motif finding.

#### III. Re-alignment of publicly available H3K27Ac dataset

Publicly available H3K27Ac ChIP-seq (*90*) data for FACS-sorted Th17 cells derived from human peripheral blood and further activated for 5 days, was acquired from GEO (GSE101389). Since the original alignment was to hg19, raw reads were obtained and re-aligned to hg38 with Burrows-Wheeler alignment (BWA). Bigwig files were generated using bam coverage, normalized to Rpkm. Input subtracted files were generated using Compare Utility from deepTools.

### SNP analysis

SNPs associated with 11 auto-immune diseases were analysed for enrichment within the TF ChIP peaks using the R package snpEnrichR (v. 0.0.1) (*159*). The SNPs were queried from the NHGRI-EBI GWAS catalogue; SNPs from studies with meta-analysis of more than one disease and from populations other than Caucasian were excluded from further analysis, and correlated SNPs were clumped (distance = 1000 kb, LD r2 = 0.8). Random SNP sets matching the disease-associated SNPs were produced using SNPsnap (*160*) server with default parameters except distance = 1000 kb, LD buddies ±20%, r2 = 0.8. Proxy SNPs for both disease-associated and random SNPs were calculated using Plink (v1.90b6.16) (https://www.cog-genomics.org/plink/1.9/) (*161*) from 1000 genomes EUR population. SNPs and their proxies (distance within 100 kb and r2 > 0.8, determined from 1000 genomes Eur population) overlapping the peaks, were identified and annotated to the nearest neighbour gene using ChIPpeakAnno. SNPs and proxies overlapping known transcription factor motifs were identified using annotatePeaks.pl from Homer. Motifs were searched within a 30-bp region around each SNP coordinate.

### Graphical representation, Venn diagrams and Statistical analysis

All graphs were plotted using GraphPad Prism software (V8.3.0). Two-tailed students T-test was used to calculate statistical significance. Venn diagrams were generated using Biovenn (*162*) or Venny (*163*). Workflow illustrations for the study were prepared used BioRender.com.

## ACKNOWLEDGMENTS

We thank all voluntary blood donors and personnel of Turku University Hospital, Department of Obstetrics and Gynecology, Maternity Ward (Hospital District of Southwest Finland) for the umbilical cord blood collection. We are grateful to Marjo Hakkarainen and Sarita Heinonen for their excellent technical help. We duly acknowledge the department’s core facilities, namely, the Finnish Functional Genomics Centre (FFGC), Cell Imaging Core (CIC) Facility and Turku Proteomics Facility supported by Biocenter Finland, for their assistance. The Finnish Centre for Scientific Computing (CSC) and ELIXIR Finland are acknowledged for computational resources.

## AUTHOR CONTRIBUTIONS

A.S designed and performed the experiments, analysed data, interpreted results, prepared figures, and wrote the manuscript; S.K.T. initiated the study, designed and performed the experiments, provided expertise, analysed data and co-wrote the manuscript; S.J performed major part of the computational analysis for the data, prepared figures and wrote part of the methods; T.B performed experiments, analysed data, prepared figures and wrote part of the methods; R.B. prepared cultures and assisted with experiments; S.D.B. analysed data and prepared figures for the protein-interaction study; T.E analysed a part of the ChIP-seq data; A.L performed preliminary analysis for the RNA-seq datasets; R.M provided expertise on the protein-interaction study; O.R. provided expertise, assisted with experiments and edited the manuscript; S.G. provided expertise and supervision and edited the manuscript; L.L.E. provided expertise, participated in the interpretation of the results, provided guidance and supervision and edited the manuscript; R.L. designed the study setup, provided expertise, participated in the interpretation of the results, provided guidance and supervision, and wrote the manuscript. All authors have contributed to the manuscript.

## DATA AVAILABILITY

The RNA-seq and ChIP-seq data reported in this paper are submitted to GEO with the accession numbers – GSE174809 and GSE174810. All other data are available in the main text or the Supplementary Materials.

## SUPPLEMENTARY INFORMATION

### SUPPLEMENTARY METHODS AND MATERIALS

#### Western blotting

Cell culture pellets were lysed using RIPA buffer (Pierce, Cat no. 89901), supplemented with protease and phosphatase inhibitors (Roche) and sonicated using Bioruptor UCD-200 (Diagenode, Seraing, Belgium). Sonicated lysates were centrifuged at 14,000 rpm for 20 min at 4°C and supernatants were collected. Samples were estimated for protein concentration (DC Protein Assay; Bio-Rad) and boiled with 6x Laemmli buffer (330 mM Tris-HCl, pH 6.8; 330 mM SDS; 6% β-ME; 170 μM bromophenol blue; 30% glycerol). Samples were loaded on gradient Mini-PROTEAN TGX Precast Protein Gels (BioRad, Helsinki, Finland) and transferred to PVDF membranes (Trans-Blot Turbo Transfer Packs, BioRad).

The following antibodies were used: anti-FOSL1 (Cell Signaling Tech, Cat no. 5281), anti-FOSL2 (Cell Signaling Tech., Cat no.19967); anti-STAT3 (Cell Signaling Tech., Cat no. 9139); anti-BATF (Cell Signaling Tech., Cat no. 8638), anti-STAT4 (Cell Signaling Tech., 2653); anti-NT5E/CD73 (Cell Signaling Tech., Cat no. 13160); anti-APOD (Santa Cruz, Cat no. sc-166612); anti-JUNB (Santa Cruz, Cat no. sc-8051); anti-RORC (eBioscience, Cat no. 14-6988-82) and anti-β-actin (SIGMA, Cat no. A5441). HRP conjugated anti-mouse IgG (SantaCruz, Cat no. sc-2005) and anti-rabbit IgG (BD Pharmingen, Cat no. 554021) were used as secondary antibodies.

#### Flow cytometry

The following antibodies were used for flow cytometry: anti-CCR6 PE (BD Cat no. 559562); anti-FOSL1 (Santacruz Biotechnology, Cat no. sc-28310); anti-FOSL2 (Cell Signaling Tech., Cat no.19967); APC-NT5E (CD73) monoclonal antibody (AD2) (Thermo Fischer, Cat no.17-0739-42), PE anti-human CD70 antibody (Biolegend, Cat no. 355103). For the primary antibodies that were unlabeled, the following secondary antibodies were used: Alexa 647 anti-mouse (Life Technologies, Cat no. A21235) and Alexa 647 anti-rabbit (Life Technologies, Cat no. A21245).

Anti-CCR6, anti-CD70 and anti-CD73 surface staining was performed 72h after initiation of Th17 culture, for which cells were washed twice with FACS buffer (0.5% FBS/0.1% Na-azide/PBS) and incubated with pre-labelled antibody for 20 min at 4°C. For intracellular staining, cultured cells (24h or 72h) were fixed and permeabilized according to the manufacturer’s instructions using IC staining buffers (Invitrogen, Cat nos. 00-5223-56; 00-5123-43; 00-8333-56). Cells were incubated with primary antibodies for 2h and subsequently washed using Perm Buffer. This was followed by 30-min incubations with labelled secondary antibodies. Suitable isotype or secondary antibody controls were maintained. Samples were acquired on LSRII (BD Biosciences, Franklin Lakes, NJ); live cells were gated based on forward and side scattering. The acquired data was analyzed with FlowJo (FLOWJO, LLC).

#### Cytokine-induction assay

Naive CD4^+^ T cells were cultured under the following conditions: CD3/CD28 activation (Th0), Th0 with IL-6, Th0 with IL-1β, Th0 with TGF-β, Th0 with IL-6 and IL-1β, Th0 with IL-6 and TGF-β, Th0 with IL-1β and TGF-β, and Th17 differentiation conditions for 24h. The neutralizing antibodies anti-IFN-γ and anti-IL-4 were added to each of these conditions. All cytokine and antibody concentrations were as described for Th17 culture conditions. FOSL1 or FOSL2 levels were estimated using Flow cytometry by performing Intracellular staining (described in the Flow Cytometry methods).

#### ELISA for IL-17 secretion

Secreted IL-17A levels were estimated using cell-culture supernatants of 72h cultured Th17 cells using either the Milliplex MAP human IL-17A kit (Merck Millipore; HCYTOMAG-60K-01), Bioplex human IL-17A Cytokine/chemokine 96-Well Plate Assay (Bio Rad; Cat. no. 171B5014M, 171304090M) or human IL-17A Duoset ELISA kit (R&D Biosystems DY317-05, DY008). The amount of IL-17A secreted by Th17 cells was normalized with the number of living cells determined based on forward and side scattering in flow cytometry analysis (LSRII flow cytometer; BD Biosciences).

#### Quantitative real-time PCR

Total RNA was isolated as described in ‘*RNA Isolation and RNA-Seq Sample Preparation’ (Main methods section)*. cDNA was synthesized using SuperScript II Reverse Transcriptase and oligo(dT) primers as described in the manufacturer’s instructions (Invitrogen, Cat nos. 18064-014 and 18418012). TaqMan primers and probes were designed with Universal Probe Library Assay Design Centre (Roche). All Taqman reactions were performed using Absolute QPCR Mix, ROX (Thermo scientific, Cat no. AB1139A). EF1α was used as endogenous control. The qPCR runs were analysed using the 7900HT Fast Real-Time PCR System (Applied Biosystems).

#### Ingenuity Pathway Analysis (IPA)

Pathway analysis was performed using Ingenuity Pathway Analysis (IPA, https://www.qiagen.com/ingenuity; Qiagen; March 2019) tool. IPA pathways with p-value < 0.01 were considered as significantly enriched.

#### Immunofluorescence analysis

CD4^+^ T cells were cultured for 72h under Th17 differentiation conditions and then spun down on poly-L-lysine-coated coverslips at 800 rpm. Cells were washed, fixed and permeabilized using Ebioscience Intracellular Staining kit (Invitrogen Cat nos.00-5223-56, 00-5123-43 and 00-8333-56). Permeabilized cells were further incubated overnight with primary antibodies against FOSL1 (Santacruz Biotechnology, Cat no. sc-28310) or FOSL2 (Cell Signaling Tech, Cat no.19967), and Lamin A/C (Santacruz Biotechnology, Cat no. sc-7292). Cells were washed with Permeabilization buffer and further incubated for 60 mins with the respective anti-mouse or anti-rabbit Alexa flour secondary antibodies (Invitrogen Cat nos. A11031; A31572; A21202). Atto-Phalloidin A647 (Sigma, Cat no. 65906) was used to stain cytoplasmic actin. Stained cells were finally mounted in Prolong Gold Antifade Mountant with DAPI (Life Technologies, Cat no. P36941) and imaged on Zeiss 780 Confocal microscope.

#### Data representation for RNA-seq and ChIP-seq data

##### I. Heatmaps

###### a. Clustered heatmap

K-means clustered heatmap was generated using the ‘PlotHeatmap’ function from DeepTools to visualize the genomic occupancy patterns of FOSL1, FOSL2 and BATF. Matrices of genomic coordinates were created using the ComputeMatrix tool, and FOSL1 peaks were used as a reference since FOSL1 ChIP-seq data analysis returned the least number of peaks amongst the three TF datasets. Genomic coordinates were further annotated using ChIPseeker (*164–166*).

###### b. Heatmap for direct targets of FOSL1, FOSL2 and BATF

Common sites for FOSL1, FOSL2 and BATF obtained from ChIP-peak Anno analysis, were annotated to the nearest TSS using Homer. Of these, the genes differentially regulated in DKD or DOE (FDR ≤ 0.1 and fold-change ≥ 1.5) were considered. Their corresponding RNA-seq expression changes were acquired using ‘Joint two files’ operation on Galaxy Europe. Subsequent heatmaps were plotted using ‘plotHeatmap’ function from deepTools (*165*).

##### II. Volcano plots for RNA-seq and ChIP-seq targets

###### a. Volcano plot for double KD, double OE and BATF KD RNA-seq targets

List of DE targets was acquired from RNA-seq analysis of double KD (24h and 72h), double OE (72h) or BATF KD (24h and 72h) Th17 cells. Volcano plots were generated using the ‘Volcano Plot’ function of Galaxy Europe (*165*). Targets with FDR ≤ 0.1 and fold change ≥ 1.8 were highlighted in red (upregulated) or blue (downregulated). Selected Th17-relevant genes were represented with labelled boxes.

###### b. Volcano plot for direct targets shared between FOSL1 and FOSL2

FOSL1 and FOSL2 common sites obtained from ChIP-peak Anno analysis, were annotated to the nearest TSS using Homer. Of these, the genes differentially regulated in double KD or double OE (FDR ≤ 0.1 and fold-change ≥ 1.5) were considered. The corresponding RNA-seq expression changes for the listed targets were acquired and subsequent volcano plots were created as described above (*165*).

#### Cytoscape network for shared interactors of FOSL1 and FOSL2

The list of shared interactors for FOSL1 and FOSL2 was obtained from a parallel study of our lab (1*67*). The common binding partners with known relevance to T-cell function were mapped against the STRING database. The assigned protein-protein interaction (PPI) network was further visualized using Cytoscape (*168*).

### STRING interactome for BATF

A predictive interactome network for BATF was acquired using the STRING database (Only ‘Text mining’ and ‘Experiments’ were considered as the information source for the predicted partners). The minimum required interaction score was set to 0.7 (high confidence). The maximum number of interactors to be displayed in first shell was restricted to 10.

### Immunoprecipitation

Immunoprecipitation for BATF was performed using Pierce MS-Compatible Magnetic IP Kit (Thermo Fischer, Cat no.90409). 72h cultured Th17 cell pellets were lysed in appropriate volumes of cell-lysis buffer provided in the kit. BATF antibody (Cell Signaling Tech., Cat no. 8638) or control rabbit IgG (Cell Signaling, Cat no. 2729) was pre-incubated with protein A/G beads for 4–5 h to form antibody-bead complexes. Lysates were first pre-cleared with control IgG-bead complexes for 3 h. The pre-cleared lysates were then incubated overnight with BATF antibody-bead complexes (test IP) or control IgG-bead complexes (negative IP control). Immunoprecipitated protein complexes were washed (following manufacturer’s protocol) and further eluted with appropriate volume of elution buffer. Eluted protein was run for immunoblotting.

Antibodies used for IP-immunoblotting are as follows: anti-BATF (Cell Signaling Tech, Cat no. 8638); anti-RUNX1 A-2 (Santa Cruz Biotechnology, Cat no. sc-365644); anti-JUNB C-11 (Santa Cruz Biotechnology, Cat no.sc-8051); anti-STAT3 (Cell Signaling Tech., Cat no. 9139); anti-IRF4 (P173) (Cell Signaling Tech., Cat no. 4964); anti-SIRT1 (Cell Signaling Tech., Cat no. 2496); anti-JUN (BD Biosciences, Cat no.610326). Conformation-specific rabbit HRP (Cell Signaling Tech., Cat no.5127) and mouse HRP (Cell Signaling Tech., Cat no. 58802) were used as secondary antibodies.

#### DNA affinity precipitation assay

DNA affinity precipitation assay experiments were performed as described in (*169–171*) with minor modifications. In brief, annealed biotinylated sense and non-biotinylated antisense bait oligonucleotides were purchased from Integrated DNA Technologies, Inc. Oligonucleotide probes containing the FOSL1, FOSL2 or BATF DNA binding motifs were designed with or without the SNP mutation. Mutations introduced to the oligonucleotides are highlighted in bold. BATF-specific and mutated sequences were used as a positive control. Neutravidin beads (Ultralink immobilized neutravidin protein, Pierce) were washed 4x with buffer A (10 mM HEPES pH 7.9, 60 mM KCl, 2 mM EDTA, 1 mM EGTA, 0.1% Triton X-100, 1 mM DTT, and protease and phosphatase inhibitors from Roche). Annealed oligonucleotides were incubated with 25 μl of beads in 200 μl buffer A for 1.5 h at 4°C with rotation at 360° rotator, followed by 4x washes with buffer A. Nuclear fraction isolated from Th17 cells cultured for 72h (Nuclear and Cytoplasmic Extraction Reagents kit, Pierce) was subjected to buffer 2 (10 mM HEPES, pH 7.9, 2 mM EDTA, 1 mM EGTA, 0.1% Triton X-100, 1 mM DTT, and protease and phosphatase inhibitors from Roche) to dilute any KCl salt. Pre-clearing was performed with unconjugated beads by incubating for 1.5 h in a 360° rotator at 4°C. Binding reactions of pre-cleared nuclear fraction with bead-conjugated oligonucleotides was performed for 4 h at 4°C, followed by washing four times with buffer A. Protein pull-down precipitates were eluted by incubating beads at 95°C for 5 min in 50 μl of 2xSDS buffer (125 mM Tris-HCl, pH 6.8, 4% w/v SDS, 20% glycerol, 100 mM DTT).

FOSL1, FOSL2 and BATF protein was analyzed by western blotting using rabbit monoclonal FOSL1 antibody (D80B4; 1:500), rabbit monoclonal FOSL2 antibody (D2F1E; 1:1000) and rabbit monoclonal BATF antibody (D7C5; 1:1000) from Cell Signaling Technology.

### Supplementary Figures and Legends

**Fig. S1.**
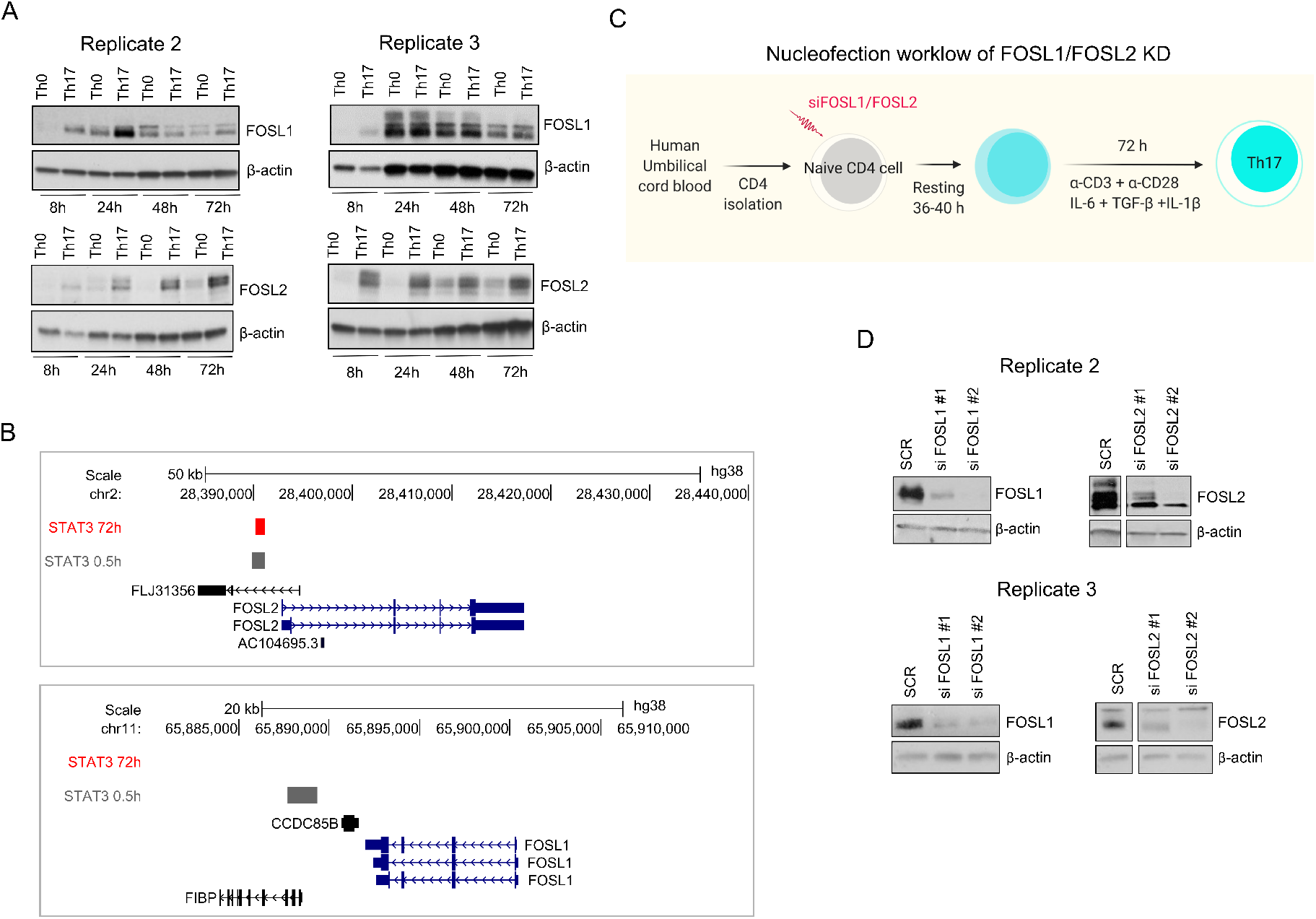
Expression and knockdown of FOSL proteins in human Th17 cells. (**A**) Immunoblot images show FOSL1 (above) and FOSL2 (below) protein levels in naive CD4^+^ T cells cultured under activation (Th0) or Th17 differentiation conditions, for the indicated time points. Actin has been used as loading control. Data represents biological replicates for Fig. 1B. (**B**) UCSC genome browser snapshots indicate the binding of STAT3 over the promoter of FOSL2 (above panel) and not FOSL1 (below panel), in Th17 cells cultured for 0.5 h and 72h. Figures were derived using bed files of STAT3 ChIP-seq data from *Tripathi et al., 2017 Cell Reports.* (**C**) Nucleofection workflow for FOSL1/FOSL2 knockdown (KD). Naive CD4^+^ T cells were treated with FOSL1- or FOSL2-targeting siRNAs, rested for 36-40 h, and further cultured under Th17-polarizing conditions (IL-6, IL-1β and TGF-β) for 72h. (**D**) Immunoblots depict FOSL1 and FOSL2 protein levels in naive CD4^+^ T cells that were silenced for the respective factors and cultured under Th17-polarizing conditions for 24h. Non-targeting siRNA (Scramble or SCR) was used as nucleofection control and actin was used as loading control. Blots shown are biological replicates for Fig. 1E.

**Fig. S2.**
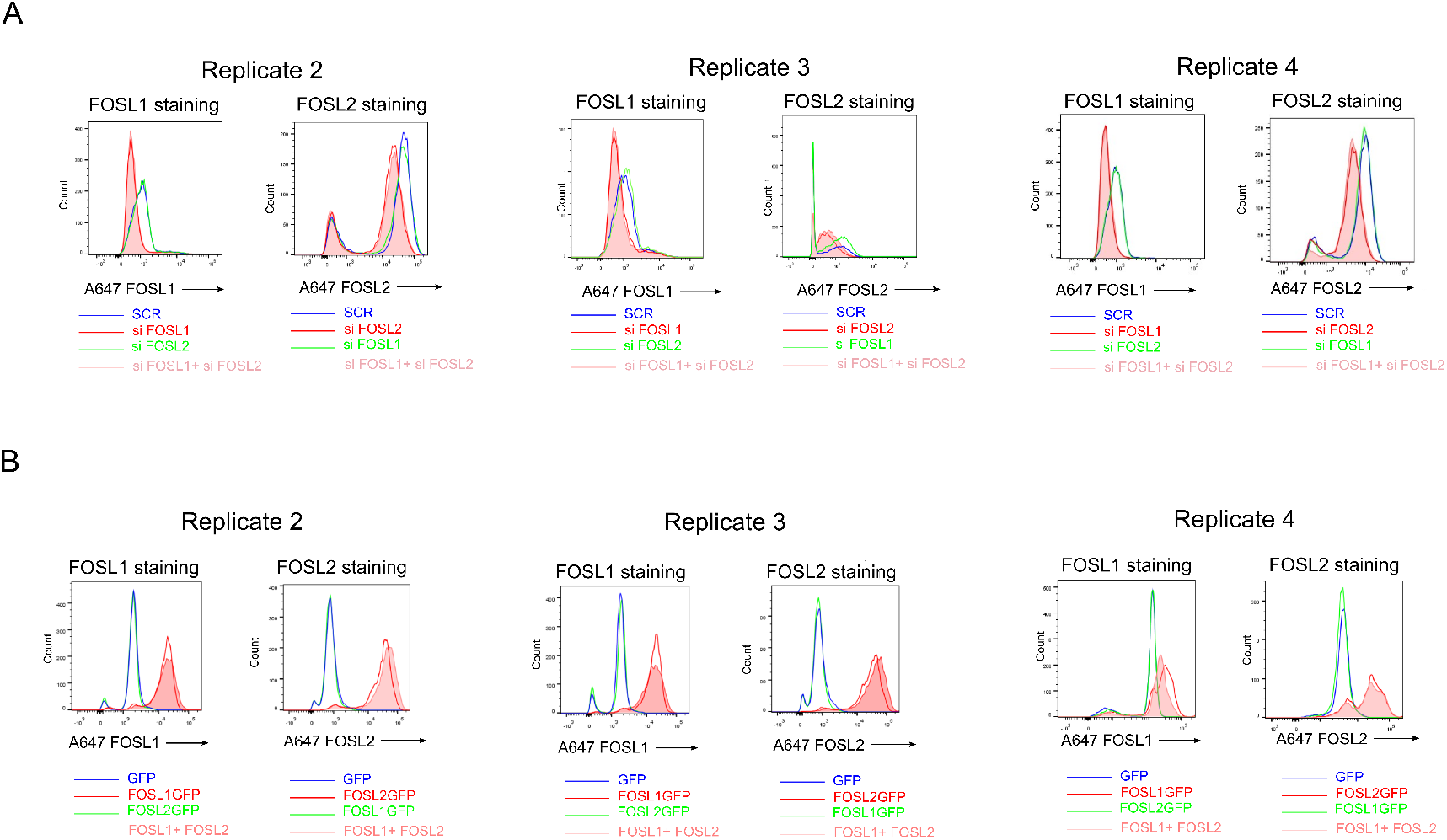
Analysis of FOSL1 and FOSL2 levels in double knockdown (DKD) and double over-expression (DOE) Th17 cells. (**A** and **B**) FOSL KD/DKD (panel A) and FOSL OE/DOE (panel B) Th17 cells were labelled (Alexa-647) for total FOSL1 and FOSL2 protein at 24h of polarization. Expression of the corresponding factors was analyzed using flow cytometry and overlay histograms were plotted [FOSL1, left; FOSL2, right]. Figure shows biological replicates for Fig. 2A and B.

**Fig. S3.**
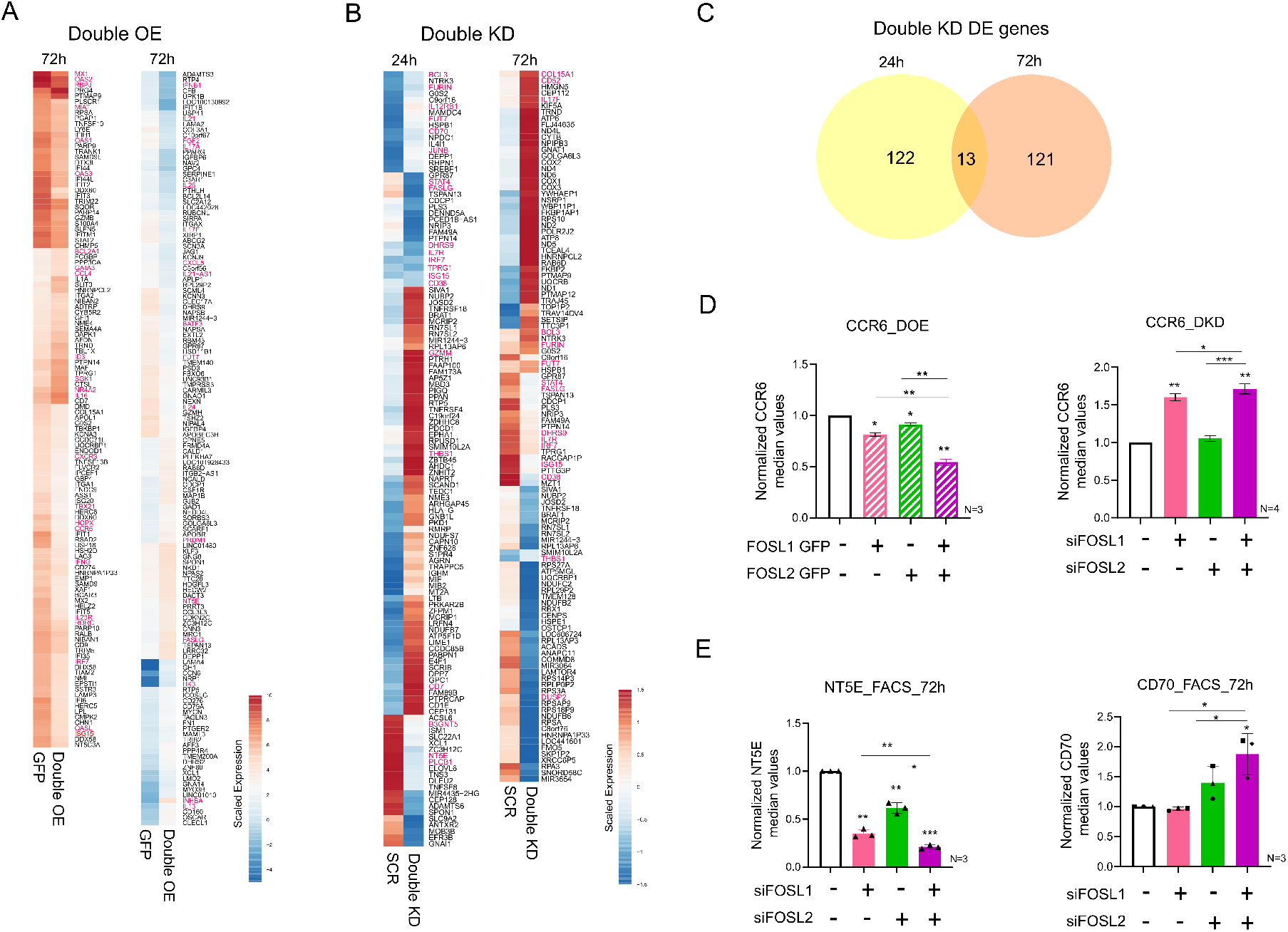
Transcriptome analysis of FOSL DKD and DOE Th17 cells and experimental validation of their targets. (**A** and **B**) Heatmap shows scaled expression values for the differentially expressed (DE) genes (FDR ≤ 0.1) in FOSL DOE (panel A) and FOSL DKD (panel B) Th17 cells. DOE targets with |fold change| ≥ 2 and DKD targets with |fold change| ≥ 1.8 are selectively shown. For the DKD heatmap, only the genes with statistically significant expression values at both time points are depicted. Targets associated with Th17-function are highlighted. (**C**) Venn diagram shows the overlap for the DKD gene-targets identified at 24h and 72h of Th17 polarization (FDR ≤ 0.1, |fold change| ≥ 1.8). (**D**) CCR6 expression was analyzed by flow-cytometry in FOSL OE/DOE [left] or KD/DKD [right] Th17 cells, at 72h of polarization. Median fluorescence intensity (MFI) values were normalized to the respective controls (SCR or Empty GFP) and plotted. (**E**) Figure shows flow-cytometry analysis of NT5E [left] and CD70 [right] protein expression in FOSL KD/DKD Th17 cells, at 72h of polarization. Bar plots show MFI values that are normalized to control. For panels D and E, data shows mean ± standard error of the mean (SEM) for three or four biological replicates, as indicated. Statistical significance was calculated using two-tailed Student’s t test (*p < 0.05, **p < 0.01, ***p < 0.001).

**Fig. S4.**
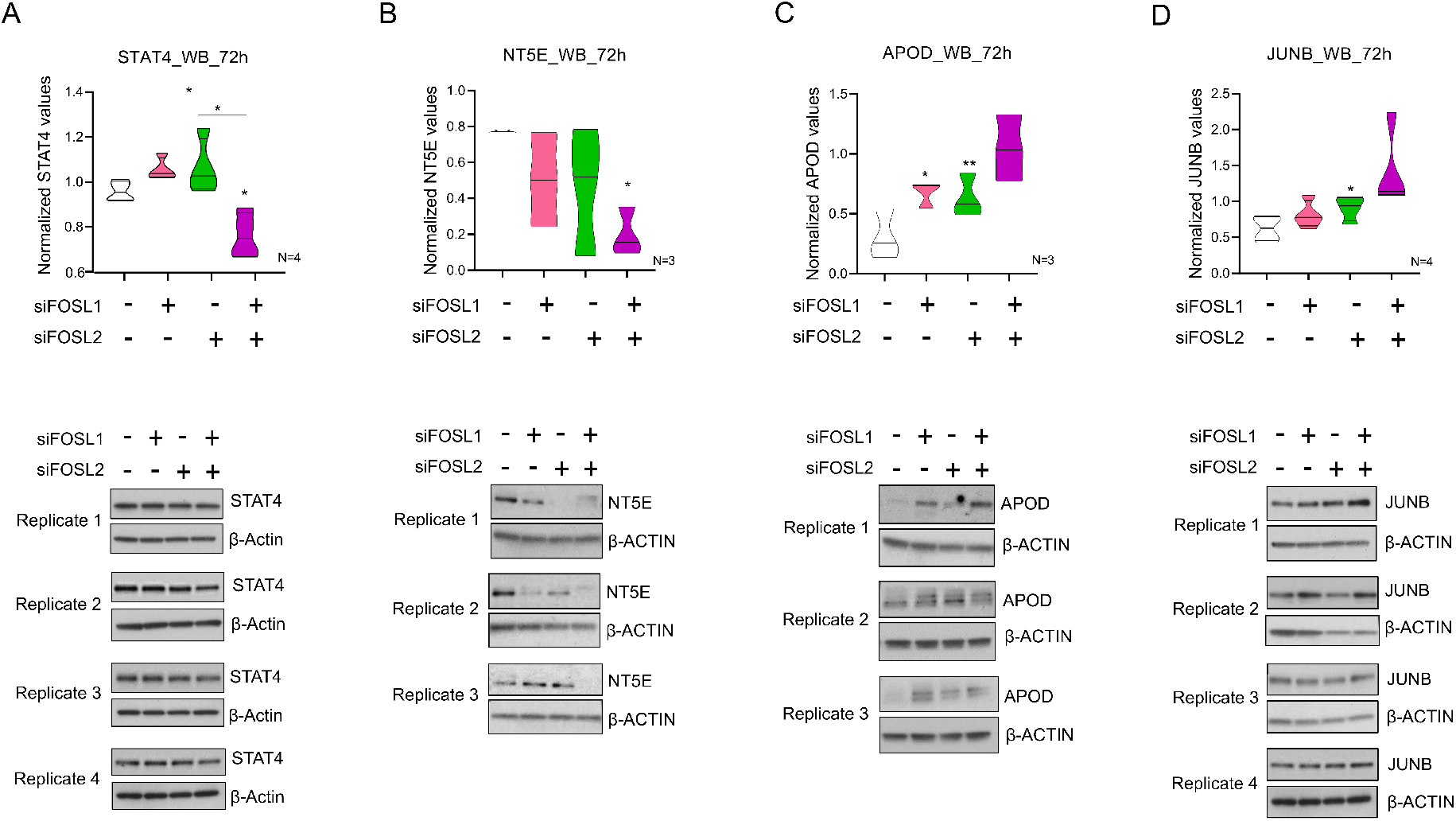
Validation of the gene-targets co-regulated by FOSL1 and FOSL2 using immunoblotting. (**A-D**) Western blots show protein-level expression of STAT4, NT5E, APOD and JUNB in FOSL KD and DKD Th17 cells at 72h of polarization. Immunoblots (below panel) were quantified using ImageJ and the corresponding FOSL intensity values were plotted into graphs (above panel). Actin served as loading control and was used for normalization. The JUNB and STAT4 blots for replicate 4 share the same loading control. Data shows mean ± SEM for three (NT5E, APOD) or four biological replicates (JUNB, STAT4). Statistical significance was calculated using two-tailed Student’s t test (*p < 0.05; **p < 0.01).

**Fig. S5.**
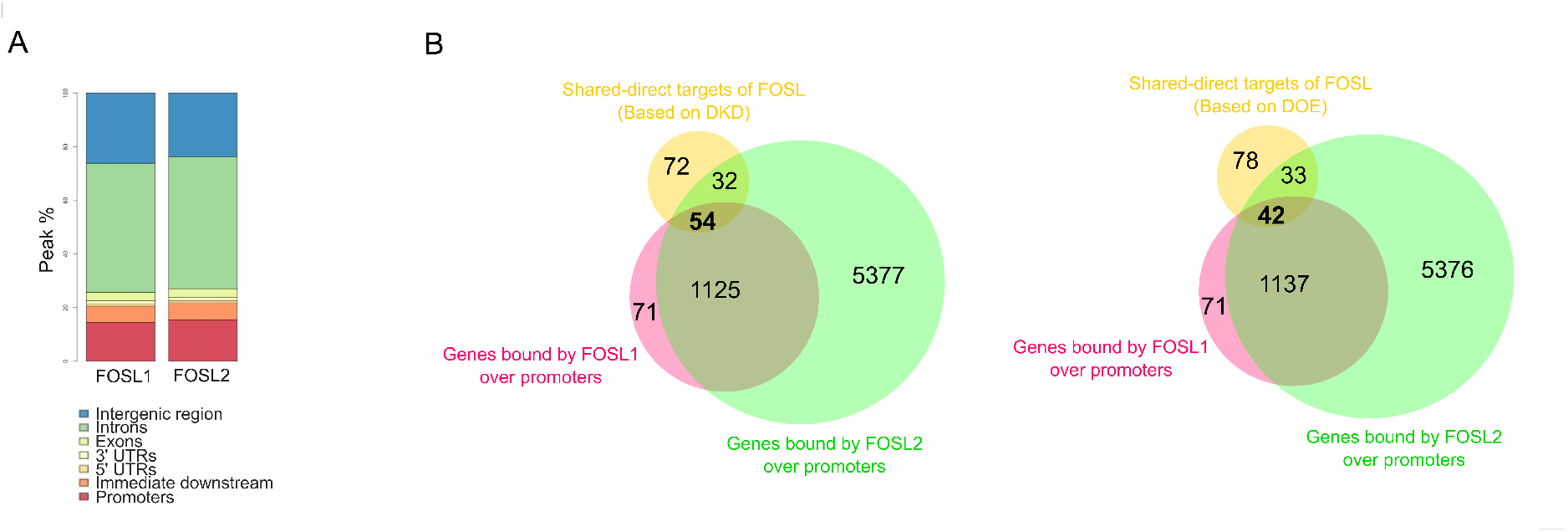
Promoter-bound shared direct targets of FOSL1 and FOSL2. (**A**) Bar plot depicts peak-annotation results for binding sites of FOSL1 and FOSL2 in 72h Th17-polarized cells. (**B**) Genes that were co-regulated (i.e., DE under DKD or DOE conditions) and showed co-localized genomic-binding of FOSL1 and FOSL2, were annotated as their shared direct targets. Venn diagram in the figure highlights (in bold) the shared targets [DKD, left; DOE, right] that are bound by FOSL factors over putative-promoter regions (5-kb around TSS). Out of these, the Th17-relevant targets have been marked in the volcano plots of Fig. 4D.

**Fig. S6.**
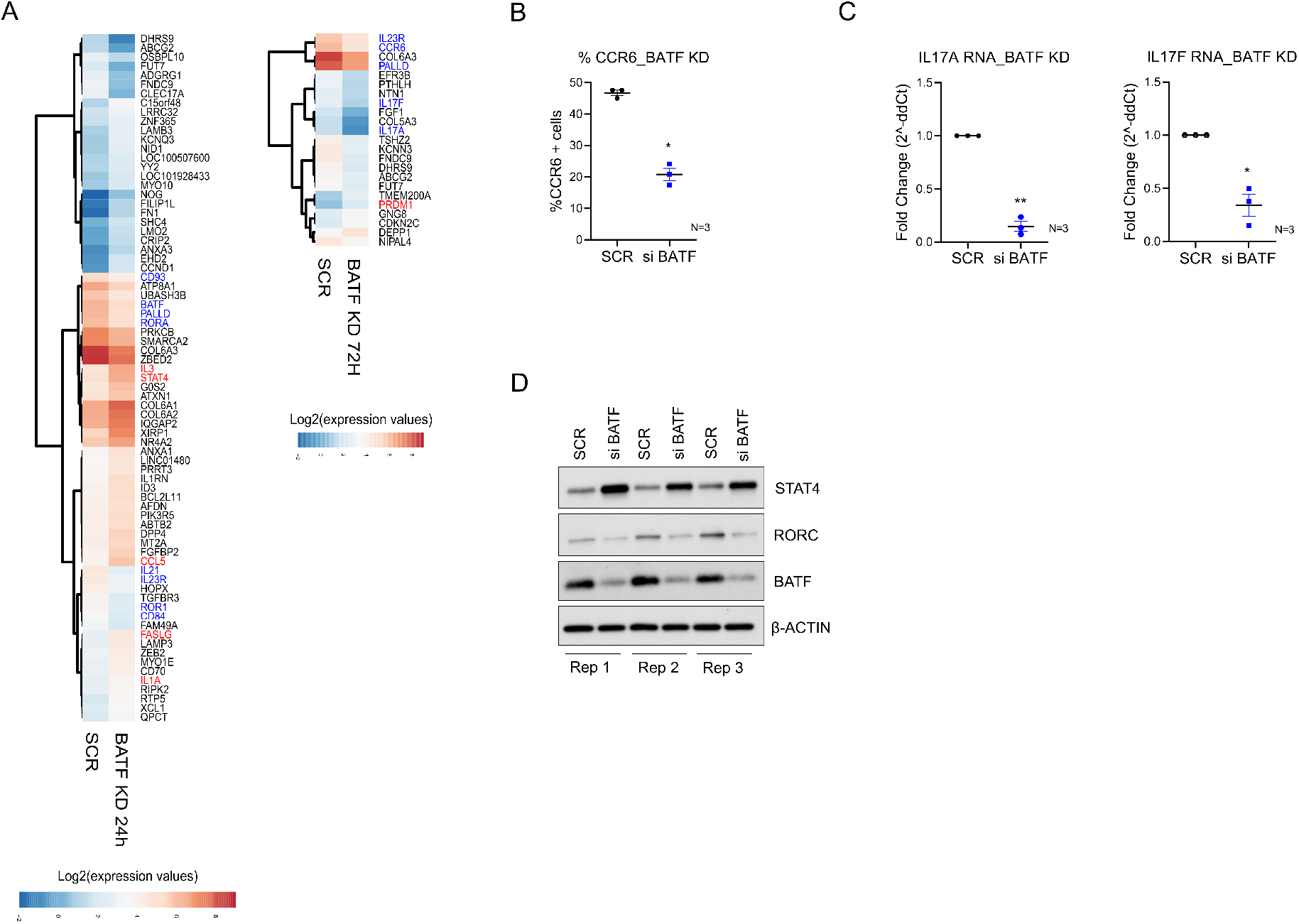
BATF gene-targets in human Th17 cells. (**A**) Heatmaps show top DE genes (FDR ≤ 0.1, |fold change| ≥ 1.8) in BATF-silenced Th17 cells at 24h [left] and 72h [right] of polarization. Scaled expression values are plotted and genes associated with Th17 cell-function are highlighted (upregulated genes are in red, downregulated genes in blue). (**B** and **C**) Panel B shows flow cytometry analysis for percentage of CCR6 positive cells in non-targeting versus BATF-silenced Th17 cells, at 72h of polarization. Panel C depicts qRT-PCR analysis for IL-17A [left] and IL-17F [right] transcript levels under the mentioned conditions. Data shows mean ± SEM for three biological replicates. Statistical significance was calculated using two-tailed Student’s t test (*p < 0.05; **p < 0.01). (**D**) Western blot analysis shows STAT4, RORC and BATF protein levels in non-targeting versus BATF KD Th17 cells, at 72h of polarization. Data for three biological replicates is shown and the quantified bar plot is provided as a part of Fig. 6F.

**Fig. S7.**
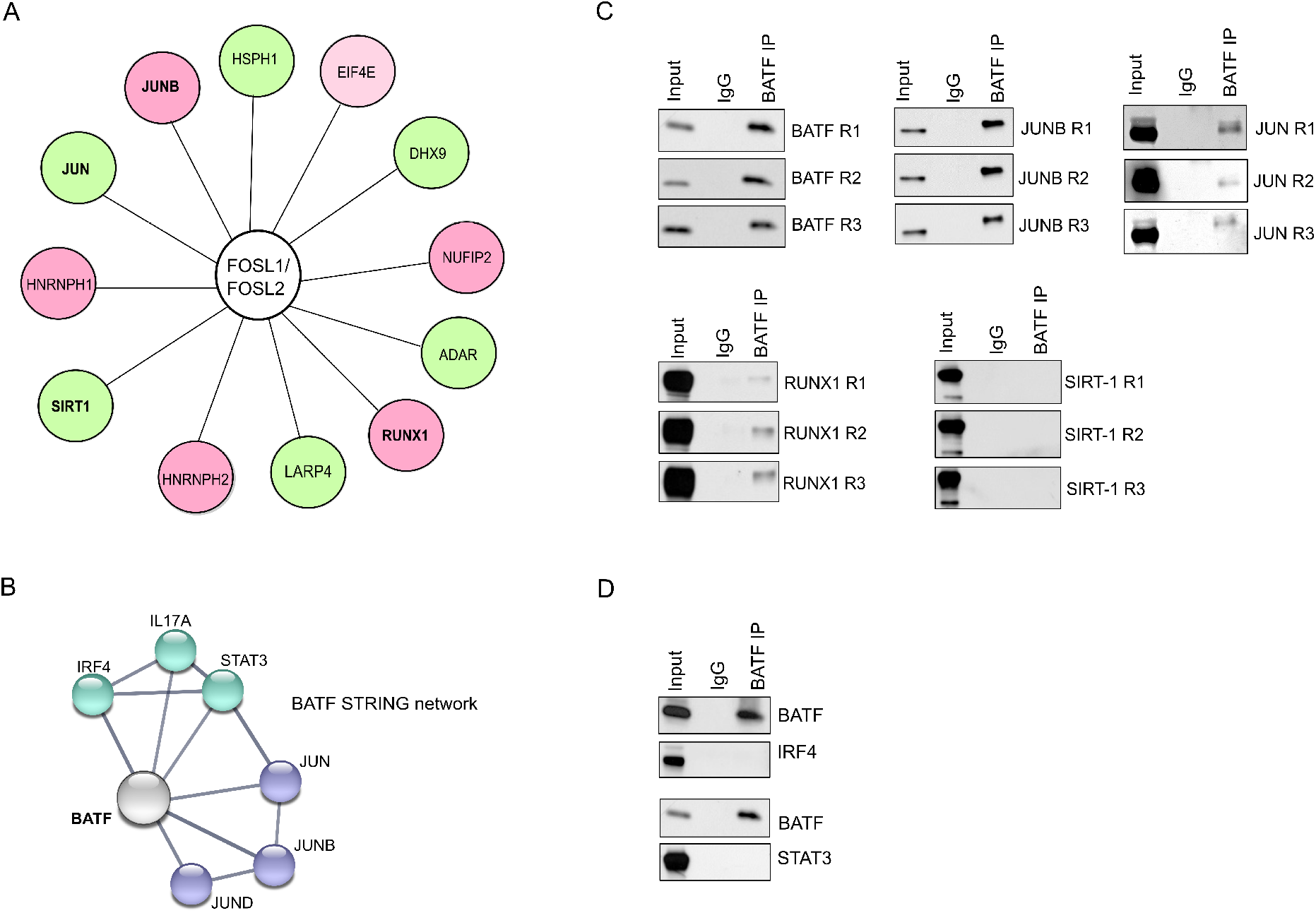
Interacting partners shared between BATF and FOSL proteins. (**A**) Figure illustrates the common binding partners of FOSL1 and FOSL2 in Th17 cells (72h), based on data acquired from a parallel study of our lab (*Shetty et al., 2021 bioRxiv*). Interactors having reported roles in T-cell function are shown. (**B**) STRING network analysis of human BATF. Width of lines between the nodes indicate confidence values for each protein-protein association. Only interactions with a minimum score of 0.7 are shown (high confidence). (**C** and **D**) Immunoprecipitated BATF was analyzed for its interaction with selected common binding partners of FOSL1 and FOSL2 (JUNB, SIRT-1, JUN and RUNX1), using western blotting (panel C). Additionally, BATF-interaction with STAT3 and IRF4 was analyzed to validate their previously-known association in mouse (panel D). Data is shown for three biological replicates.

**Fig. S8.**
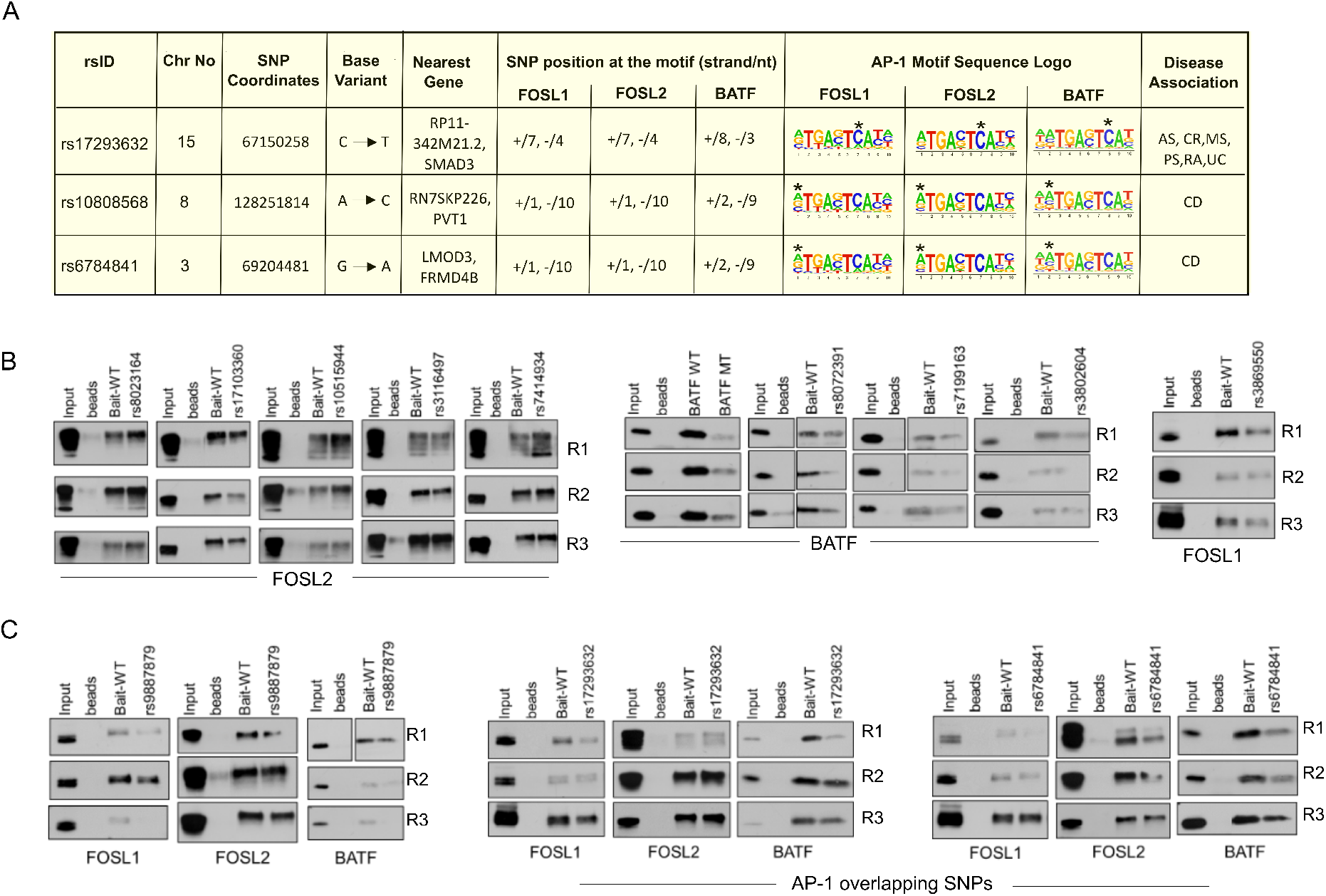
Disease-linked SNPs overlapping with AP-1 motifs and DAPA analysis of selected SNPs. (**A**) Table illustrates information on the autoimmune-linked SNPs that are harbored within consensus AP-1 motifs at the shared genomic-binding sites of FOSL1, FOSL2 and BATF. The sequence logos shown have been derived from the respective TF ChIP-seq peaks using Homer. (**B** and **C**) DAPA analysis was performed to test the effect of selected SNPs on the DNA-binding abilities of FOSL1, FOSL2 and BATF. The immunoblot images in panels B & C show biological replicates (R1, R2, R3) for Fig. 7C and Fig. 7D respectively

